# Aggression and multimodal signaling in noise in a common urban songbird

**DOI:** 10.1101/2022.04.29.490020

**Authors:** Çağla Önsal, Alper Yelimlieş, Çağlar Akçay

## Abstract

Anthropogenic noise may disrupt signals used to mediate aggressive interactions, leading to more physical aggression between opponents. One solution to this problem is to switch signaling effort to a less noisy modality (e.g., the visual modality). In the present study we investigate aggressive behaviors and signaling in urban and rural male European robins (*Erithacus rubecula*) in response to simulated intrusions with or without experimental noise. First, we predicted that urban birds, living in noisier habitats, would be generally more aggressive than rural birds. We also predicted that during simulated intrusions with experimental noise, robins would increase their physical aggression and show a multi-modal shift, i.e., respond with more visual threat displays and sing fewer songs. Finally, we expected the multi-modal shift in response to noise to be stronger in urban birds compared to rural birds. The results showed that urban birds were more aggressive than rural robins, but an increase in aggression with experimental noise was seen only in the rural birds. Urban but not rural birds decreased their song rate in response to noise. Contrary to the multi-modal shift hypothesis, however, there was no evidence of a concurrent increase in visual signals. These results point to a complex role of immediate plasticity and longer-term processes in affecting communication during aggressive interactions under anthropogenic noise.

**Significance Statement:** Human activity has an enormous effect on wildlife, including on their social behavior. Animals living in urban areas often tend to be more aggressive than those living in rural areas, which may be due to urban acoustic noise making communication between individuals more difficult. In a study with a common songbird, the European robin, we investigated the role of urban acoustic noise in aggression and territorial communication. Urban robins were more aggressive than rural robins, and additional noise in the territory increased aggression in rural but not urban robins. While urban robins decreased their singing effort with additional noise, they did not increase visual signals concurrently. These results suggest that noise can indeed make animals behave more aggressively although the effect may depend on how noisy it is already. These results further our understanding of how human-made noise changes animal communication and social behavior.

## Introduction

Urban habitats are polluted with anthropogenic noise, often in multiple modalities, which creates challenges for urban-living wildlife (Brumm and Slabbekoorn 2005). Many species rely heavily on signals for communication in contexts such as mate attraction and territorial defense, and noise from vehicles, buildings and other human activities often interferes with these signals (Francis et al. 2009; Halfwerk and Slabbekoorn 2015; Lee and Thornton 2021). A well-studied example of the effect of anthropogenic noise on communication is vocal signaling in urban birds: in response to anthropogenic noise commonly found in cities, many species of birds may increase repetition rates, amplitude, or frequency characteristics of their acoustic signals (Brumm 2004; Gil and Brumm 2014; Slabbekoorn and den Boer-Visser 2006; Wood and Yezerinac 2006).

Urban living also leads to increased aggressiveness of individuals in urban habitats compared to the rural habitats (Davies and Sewall 2016; Evans et al. 2010; Hardman and Dalesman 2018; Phillips and Derryberry 2018; Scales et al 2011). The reasons for increased aggression in urban habitat are not yet fully understood. It may result from several factors, including selection for bolder individuals (Evans et al. 2010), increased food resources (Foltz et al. 2015), increased exposure to harmful chemicals such as lead (McClelland et al. 2019) and less stable social environment due to high rates of territory turnover in urban habitats (Davis et al. 2013). Anthropogenic noise in urban habitats may also be responsible for increased aggression.

Animals often use signals in aggressive interactions (e.g., during territory defense) to resolve conflicts with opponents. Use of signals is often beneficial for both parties if they can avoid costly physical fights in this way (Maynard Smith and Price 1973). Consequently, if signaling is prevented or the signals are rendered ineffective, individuals may need to resort to higher levels of physical aggression (Logue et al. 2010). Applied to urban habitats, this hypothesis suggests that the high levels of urban noise may render long-distance aggressive signals less effective, which in turn may lead to higher levels of aggression (Phillips and Derryberry 2018). Consistent with this hypothesis, some studies reported a positive correlation between ambient noise levels and aggressive behaviors (Akçay et al. 2020; Phillips and Derryberry 2018; but see Kleist et al. 2016).

Signalers employ various strategies to overcome interference from anthropogenic noise. We focus here on the flexibility afforded by having signals in more than one modality (Bro-Jørgensen 2010; Halfwerk and Slabbekoorn 2015; Partan and Marler 1999). When animals have signals in more than one modality, they may shift their signaling effort from the noisy modality to a less noisy modality to increase the likelihood that the message of the signals gets through to the receivers (Partan et al. 2010; Partan 2017). This hypothesis is termed the multi-modal shift hypothesis.

Few studies tested the multi-modal shift hypothesis in signals used in territorial interactions. In one study, Ríos-Chelén et al. (2015) found that male red-winged blackbirds (*Agelaius phoeniceus*) did not use more intense visual signaling in noisier territories although they modified their acoustic signals (e.g., decreased their song rate). Another study on song sparrows (*Melospiza melodia*) found that males in noisier urban habitats were both more aggressive and used proportionally more visual threat signals (wing waves) during territory defense compared to the males in rural habitats, consistent with a multi-modal shift (Akçay et al. 2020). In a further experiment, however, individual song sparrows did not increase their visual signaling effort when experimentally presented with noise, suggesting that multi-modal shift seen in urban song sparrows was not due to immediate plasticity (Akçay and Beecher 2019).

Here we investigate the responses of European robins (*Erithacus rubecula*) living in urban and rural habitats in Istanbul, Turkey, to simulated territorial intrusions with or without experimental noise playback. European robins have both visual and acoustic signals that are used in agonistic interactions (Lack 1943). Previous studies have found that robins respond both to song and visual signals during territorial intrusions (Chantrey and Workman 1984). Territory holders sing in response to the song of an intruder, while visual signals are used when the intruding male is within range of vision. Their most prominent visual signal is the neck display, which has been observed in response to the sight of a rival male’s red neck, or indeed even a ball of red feathers (Lack 1943). Other visual threat signals include wing flutters, pricking the tail up, and swaying, where the resident male moves his head from one side to the other (Lack 1943).

Signaling and aggressive behaviors of robins have been investigated in multiple studies. A study by Mclaughlin and Kunc (2013) found that robins, after being lured by playback of a robin song from a speaker, tended to move away from the speaker when the speaker switched to playback of low-frequency noise mimicking typical traffic noise, particularly when the amplitude of noise was high (90 dB at 1m). In response increasing amplitude of noise, the robins sang shorter songs with fewer notes and increased the minimum frequency of their songs. In another study, experimental presentation of wind turbine noise during simulated territorial intrusions led to a decrease in low frequency elements in the songs, at the same time leading to an increase in flight rates (Zwart et al. 2016). Song rates did not significantly differ between the noise and no-noise treatments. Interestingly, fewer robins used visual threat postures under experimental presentation of noise compared to no noise, although the difference was not significant (Zwart et al. 2016).

The presence of both acoustic and visual signals in territorial defense makes the European robin a suitable candidate for testing the multimodal shift hypothesis but to our knowledge no previous study compared visual signaling between urban and rural robins. We predict that robins in urban habitats will exhibit higher levels of visual signaling. Additionally, if such a multi-modal shift is due to phenotypic plasticity, robins should increase their visual signaling under experimental noise. We also predicted that urban birds would exhibit higher levels of plasticity in signaling than rural birds, because they more frequently experience significant anthropogenic noise levels (Gentry et al. 2017; Lazerte et al. 2016). Finally, in accordance with earlier studies, we also expect to see a greater level of aggression from urban robins compared to rural robins. If increased aggression is due to individual plasticity in response to noise, we also expect higher levels of aggression from robins in response to experimental noise, particularly in urban birds who have more experience with noise.

## Methods

### *Study site*s and species

We carried out playback experiments with male European robins that held territories in rural areas (forests around Sarıyer, Istanbul, 41° 9’ 50.73971”N, 29° 0’ 32.25243”E) and urban parks and green areas in Sarıyer, Istanbul, Turkey in April and May 2021 (urban: n=9; rural: n=12). Robin territories were detected by the presence of an already-singing male robin before the first playback or during recording sessions prior to playback. We determined a central location by observing the robin’s flights for about 5 minutes, although we did not attempt to map the entire territory. In all trials reported below, only a single bird responded to the playback (for one subject, not included in the final sample, we aborted the trial when a second male came to within 10 m of the speaker). It was not possible to record data blind as our study involved observing focal individuals in urban or rural habitats and noise manipulation was audible to all observers.

### Stimuli

Playback stimuli were generated on the software Syrinx (John Burt, Portland, OR) from male European robin songs recorded in March 2021 in four of the nine study sites. We generated stimulus files by extracting high quality songs from each recording and filtering out low frequency noise below 1000 Hz. We added a silent period after each song so that stimuli were presented at a rate of one song per seven seconds. The songs lasted on average (±SD) 2.36 (±0.49). We created one-minute stimuli (consisting of nine different songs) which were repeated three times to make up three-minute stimuli to be played during the trials. In total, we used 17 files created from the songs of 17 different robins. The stimuli played for each subject came from a robin whose territory was separated by at least one km from that of the subject’s territory. Subjects received the same song stimulus in both trials. As a visual stimulus, we used a 3-D printed bird model (dimensions, height: 8 cm, length: 12 cm, width: 4.5 cm) which was hand-painted to resemble an adult robin (Figure S6).

We generated the experimental acoustic noise stimuli by filtering white noise (created with Audacity) with the average amplitude spectrum from a 1-minute recording (made with a Marantz PMD660 and ME66/K6 microphone) of traffic noise in Sarıyer, Istanbul using the package *seewave* in R (Sueur et al. 2008). The power spectrum of the noise stimulus can be found in the supplementary materials (Figure S1).

### Experimental procedure and design

We started each trial by placing the robin model attached to a speaker (Anker Soundcore Bluetooth Speaker, Anker, Inc.) on a natural perch at the estimated center of the resident male’s territory, approximately 1.5 m above the ground. A second Bluetooth speaker (same model as above) was placed on the ground, face-up below the first, for noise playback. In the control treatment, the second speaker was placed but not turned on, so the resident male received only song playback. In the noise treatment, in addition to the song playback, traffic noise was played at 75 dB SPL at 1 m. The noise playback lasted for the entire duration of the song playback. This stimulus simulates an acute, if transient, source of noise (e.g., a vehicle idling, or a lawnmower operating).

Each subject received two 3-minute trials, one with experimental noise and one without noise. The order of the treatments was counterbalanced, and the two trials were separated by approximately an hour. Two observers, about 10 m away from the experimental setup, recorded the songs and calls of the resident robin. The observers also narrated the trials, verbally noting flights (any airborne movement by the bird), distance with each flight, and visual displays described in Table 1 onto the recording. We continued recording songs for 3 minutes after the end of each trial. Recordings were made on a Marantz PMD660 with a Sennheiser ME66/K6 microphone, or on a Zoom H5 handheld recorder with a Zoom SGH6 shotgun microphone.

**Table 1.** Visual displays of European robins during territorial interactions

| Behavior | Description |
| --- | --- |
| Neck display | The robin raises his head, displaying his neck. |
| Wing flutter | The robin flutters his wings. |
| Swaying | The robin rhythmically sways his body from one side to the other. |
| Tail up | The robin perks his tail up. |

In 31 of the trials, the bird was already singing when we started the trial. For these birds, the 3-minute playback period started with the first song played. In 11 of the trials, where the subject was quiet when the playback started, the 3-minute trial period began with their first response (song or approach). The average duration of pretrial playback for these 11 trials was 64.9 seconds (SD = 40.8).

After each trial, we measured the ambient noise with a VLIKE VL6708 sound-level meter with the method described in (Brumm 2004). We took eight measurements (two in each cardinal direction) within a minute period, which were then averaged. For three subjects, we only had noise measurements from a single trial.

### Response variables

We scanned and annotated the narrated trial recordings using the Syrinx software. The number of songs and visual displays (number of neck displays and wing flutters as well as the start and end times for swaying in seconds) were extracted from the verbal notes made on the recordings. We only analyzed song rates and durations, as overlapping stimulus and subject songs made it impossible to determine with certainty the note compositions for most songs, precluding frequency measurements. Only 8 subjects used any visual displays during the experimental period, we therefore coded visual displays as a binomial variable (visual signal present vs. absent during a trial). From the recordings, we also extracted the number of flights, closest approach to the model/speaker and proportion of time spent within 5m of the speaker. These three spatial variables were taken as aggressive behaviors.

Because the spatial variables of aggression were significantly correlated with each other (all p< 0.05), we carried out a principal component analysis (PCA) using the *principal* function in package *psych* (Revelle 2021). The first component of PCA (PCA1) explained 61% of variance and was taken as our primary measure of aggression (see Table 2 for loading coefficients). The aggression scores thus calculated have been shown to be valid measures of territorial aggression in songbirds (Akçay et al. 2013). We report the analyses using raw spatial measures in the supplementary materials.

**Table 2.**
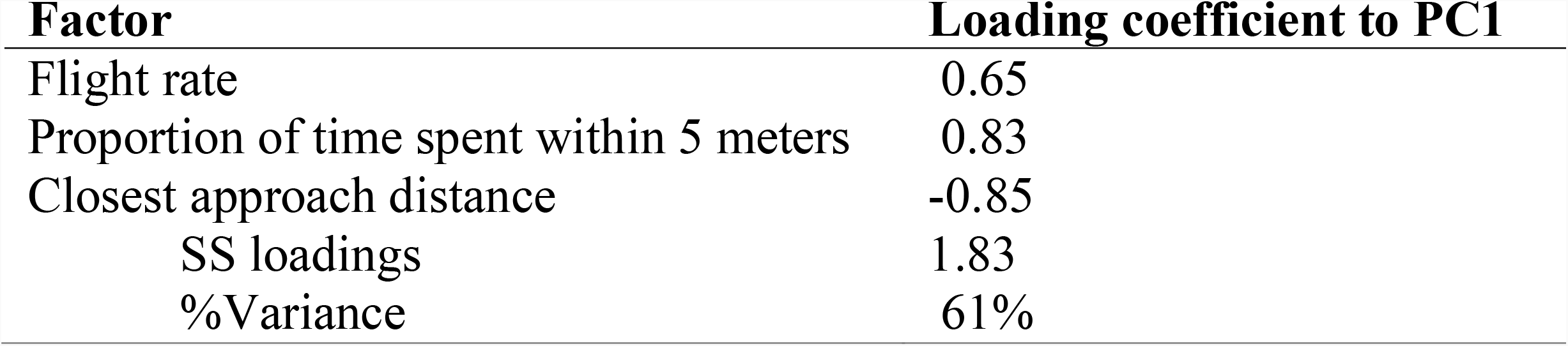
Loading coefficients of the Principal Component Analysis.

### Data analysis

All analyses were carried out in R (R Core Team 2021). We first checked whether urban territories had higher ambient noise with a linear mixed model (LMM) using habitat type (urban vs. rural) as the predictor variable and territory ID as the random variable. We also assessed whether noise levels were repeatable using the *rptR* package (Stoffel et al. 2017).

We then checked whether the order of trials had a significant effect on aggression scores, song rates and visual signaling. The order of trials did not have a significant effect on song rate (LMM, coefficient = -0.68, SE = 0.60, p = 0.28) or aggression score (LMM, coefficient = -0.22, SE = 0.15, p = 0.16). However, there was a significant order effect on the incidence of visual displays (GLMM, estimate = -15.32, SE = 6.22, p = 0.01). Eight subjects used visual displays in the first trial, compared to two in the second trial (both subjects also used visual displays in the first trial). Because of this order effect, we only used the first trial for each subject in models including visual displays.

We analyzed song rates and aggression scores with linear mixed models (LMM), using the *lme* function in the package *nlme* (Pinheiro et al. 2022). We took habitat type (urban vs. rural) and experimental treatment (noise vs. control), and their interaction as the predictor variables, and male ID as the random variable. We applied a generalized linear model (with log-link, using the “*glm*” function in base R) with visual displays as binomial response variable, and habitat and treatment as predictor variables, using only the first trials for each subject. Since only two rural subjects used visual displays, we also carried out this analysis with the subset of only urban birds (see Supplementary Materials).

## Results

Urban habitats had significantly higher levels of ambient noise than rural habitats (urban: M = 49.0, SD = 7.1; rural: M = 39.9, SD = 3.6) and noise measurements were highly repeatable between the two trials (intra-class correlation coefficient; r=0.96, standard error: 0.02; p < 0.0001).

Urban birds were significantly more aggressive than rural birds. There was no main effect of noise treatment but there was a significant interaction effect of habitat and noise treatment (Table 3, Figure 1a). To understand this interaction effect, we carried out paired t-tests on rural and urban birds with noise treatment as predictor variable. Rural birds were more aggressive under the noise compared to no noise treatment (paired t-test; t(11)=2.44, p= 0.033), whereas there was no effect of experimental noise on aggression in urban birds (paired t-test: t(8)=-1.23, p= 0.25; Figure 1a). Looking at the spatial variables separately, this interaction effect seems to be driven mostly by the closest approach measure (see Supplementary Materials).

**Table 3.** Coefficients (SE) from the linear mixed models and the p-values from Wald t tests, examining the effect of habitat and experimental noise treatment. Statistically significant values are shown in bold type

| *Predictors* | **Aggression Score** |  | **Song Rate** |  | **Song Length** |  |
| --- | --- | --- | --- | --- | --- | --- |
|  | *Estimates (SE)* | *p* | *Estimates (SE)* | *p* | *Estimates (SE)* | *p* |
| **(Intercept)** | -0.58 (0.26) | **0.035** | 8.44 (0.59) | **<0.001** | 1.86(0.14) | **<0.001** |
| **Treatment** | 1.32 (0.39) | **0.003** | 0.61 (0.65) | 0.36 | 0.04(0.21) | 0.8454 |
| **Habitat** | 0.32 (0.19) | 0.105 | 0.81 (0.90) | 0.38 | 0.03(0.11) | 0.7890 |
| **Treatment\*Habitat** | -0.68 (0.29) | **0.031** | -2.02 (1.00) | 0.059 | 0.11(0.17) | 0.5376 |

**Figure 1.**
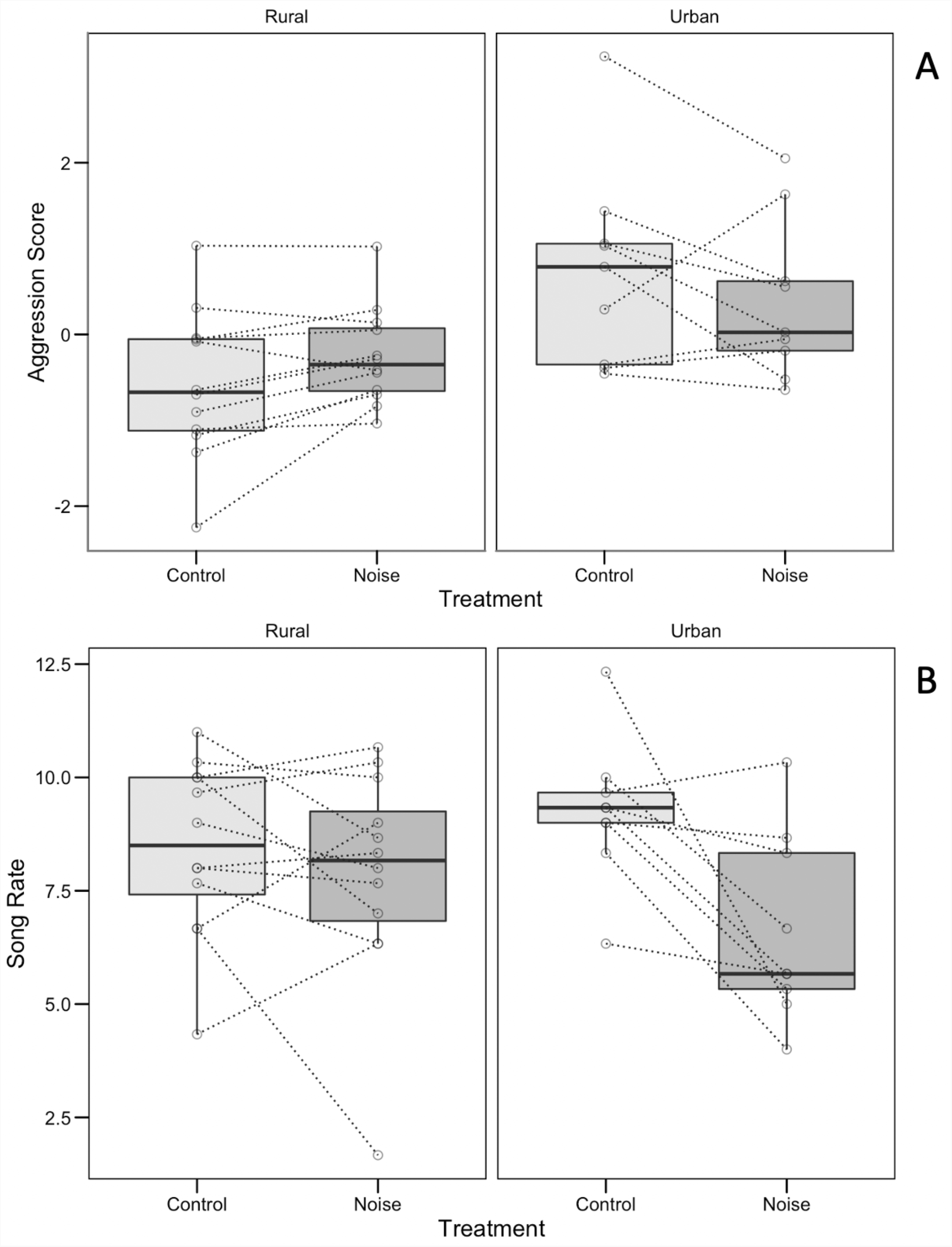
The relationship of aggression scores (A) and song rate (B) with habitat and noise treatment. The boxes indicate interquartile ranges, the middle line indicates median, and whiskers indicate 95% confidence intervals. Dots connected by dotted lines represent data from individual subjects.

Song rates did not differ significantly between urban and rural birds and the noise treatment had no main effect. The interaction effect of habitat and noise treatment however approached significance (Table 3, Figure 1b). When we analyzed song rates for urban and rural birds separately there was a significant effect of noise treatment in urban birds with lower song rates under experimental noise compared to without noise (paired t-test; t(9)=3.15, p= 0.014); while there was no effect of noise treatment for rural birds (paired t-test; t(11)=1.01, p= 0.33; Figure 1b). There was no significant main or interaction effect of habitat or noise treatment on song length (Table 3).

Urban birds used visual threat displays in the first trials significantly more than their rural counterparts (GLM; χ^2^= 9.75, p=0.0018; Figure 2). While there was no main effect of noise treatment (χ^2^= 0.00, p=1.0), the interaction effect of treatment and habitat approached significance (χ^2^= 3.72, p=0.053). This interaction effect was driven by a tendency in urban birds to use more visual signals in the no noise treatment compared to noise treatment, although the effect was not significant (GLM within the subset of urban birds; χ^2^= 3.22, p=0.07). Out of the five urban birds that received the no noise treatment in the first trial, all five used visual signals, compared to only one out of the 4 birds which received the noise treatment first (Figure 2).

**Figure 2.**
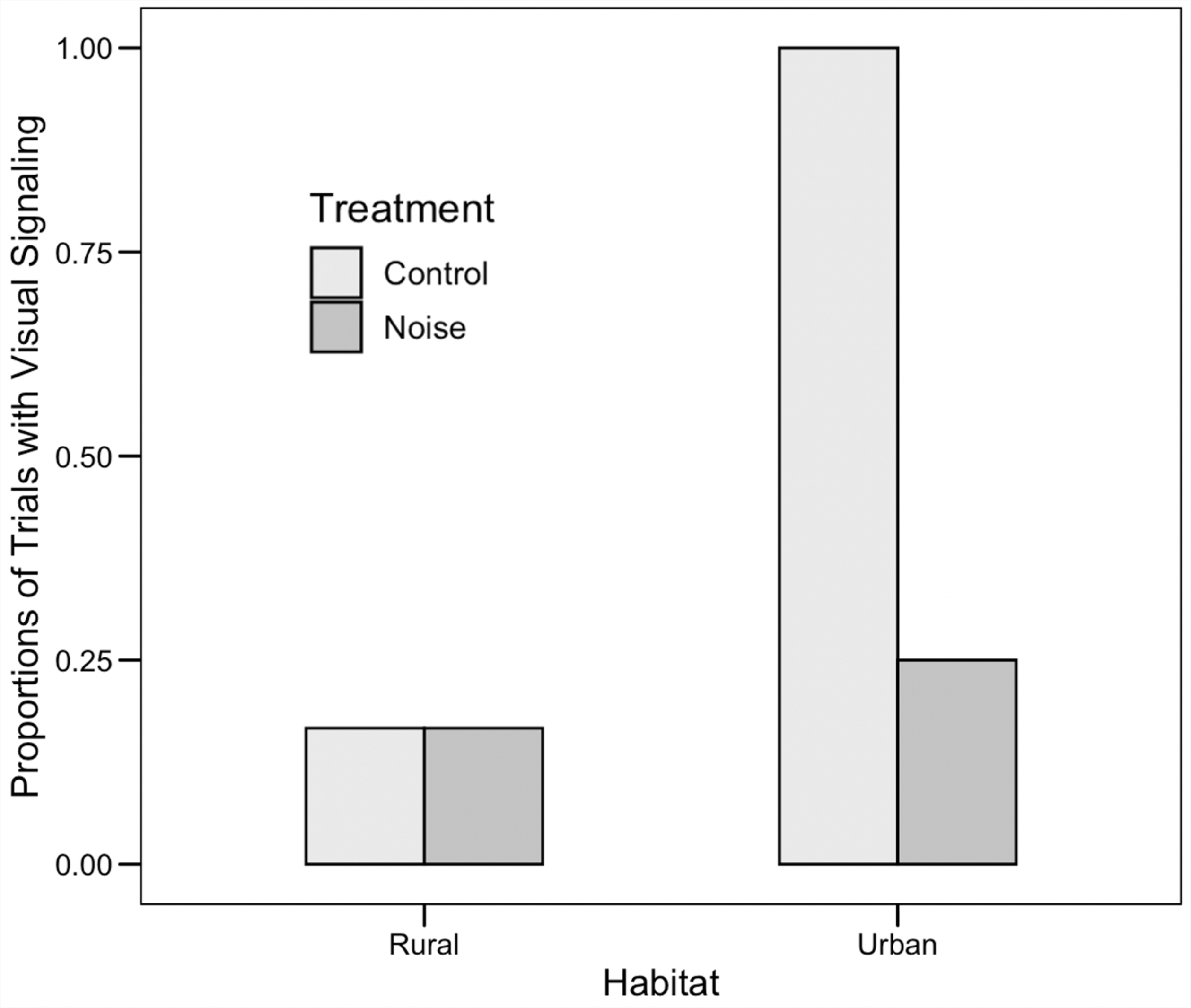
Proportion of first trials where the resident male used visual signals, grouped by habitat and noise treatment. The numbers at the bottom of each bar indicates the total number of subjects for each combination.

## Discussion

In the present study, we examined the role of acute anthropogenic noise in determining aggressiveness and aggressive signaling. We predicted that urban robins, living in noisier territories, would be more aggressive compared to rural robins in simulated territorial intrusions and experimental noise during simulated intrusions will change both their aggressive behaviors and signaling behaviors. Particularly, we expected that experimental acoustic noise should increase aggression during intrusions and lead to an increase in using signals in the visual modality, particularly in urban birds who would have more experience dealing with fluctuating levels of noise.

In line with our first prediction, we found that urban robins responded with significantly more aggressive behaviors (particularly close approach) to simulated intrusions than rural robins. The effect of experimental noise treatment on aggressive approach was dependent on the habitat: contrary to our hypothesis, experimental noise led to increased aggression in the (comparatively quiet) rural habitats, but it had no effect in the noisy urban habitats. Experimental noise led to a decrease in song rates in urban and not rural birds. No change was observed in song duration (see Table 3). Finally, visual signals were more common in urban habitats (consistent with the fact that urban birds are more aggressive) and tended to be less common under experimental noise in the urban habitats, albeit not significantly.

### Noise and aggressive behaviors in territory defense

Our results on the effect of habitat on aggression replicates earlier findings that urban-living birds are more aggressive than rural birds (Davies and Sewall 2016; Evans et al. 2010; Hardman and Dalesman 2018; Phillips and Derryberry 2018). We also extend previous findings by showing that the effect of experimental noise on aggression was dependent on habitat: urban males showed no further increases in aggression with experimental noise while rural males showed a significant increase in aggression. This is opposite of our expectation that urban birds would show higher levels of phenotypic plasticity in response to experimental noise treatment (Gentry et al. 2017; LaZerte et al. 2016). The lack of an effect of noise on aggressive behaviors in urban habitats may be due to several reasons: First, for urban males that are already living in noisy territories, additional noise may not have as much as an effect as in rural habitats. Urban birds may also be more habituated to acute increases of noise than rural birds, although they did show a plastic response in their singing rate as discussed below. Finally, urban birds may not be able to increase their already high levels of aggression in response to noise playback.

The increased aggression with experimental noise in rural habitats is consistent with the idea that urban noise has a causal role in increasing aggression. Only a small number of studies experimentally manipulated noise levels to examine a causal role of noise in increased aggression. These studies yielded mixed results. Grabarcyzk and Gill (2019) found that house wrens (*Troglodytes aedon*) males attacked the simulated intruder more frequently when playback was accompanied with experimental noise than when it wasn’t, consistent with the hypothesis that noise induces higher levels of aggression. Another study in song sparrows however, found no effect of experimental noise on aggression levels, measured as time spent within one meter of the speaker, or attacks (Akçay and Beecher 2019). In the latter study, the noise playback started only when subjects approached to within five meters, which all subjects did within a short period of time (< 1 minute). Thus, lack of an effect in physical proximity may be due the fact that subjects already were close to the speaker when the noise playback started.

In another study, Zwart et al. (2016) found that European robins did not show a statistically significant increase in aggressive behaviors in response to experimental wind turbine noise during simulated intrusions, although some variables like flights did show a trend consistent with higher aggression with experimental noise. A more recent study by Reed et al (2021) in lazuli buntings (*Passerina amoena*) and spotted towhees (*Pipilo maculatus*) found that experimental presentation of natural noise (such as noise from a river, ocean surf or cicadas) at the landscape level led to slower detection of a simulated intruder and consequently weaker approach responses (see also Kleist et al. 2016). Finally, a study conducted with saffron finches (*Sicalis flaveola*) found birds displayed lower agonistic behaviors under experimental traffic noise, although this study is harder to interpret in the present context as the experiment was done with captive birds that lived in small cages (Passos et al. 2020).

Together these studies point to two apparently contradictory effects of noise on territorial aggression. On one hand, noise may make localization of the simulated intruder and perception of stimulus features more difficult, leading to slower or weaker approach behaviors (Kleist et al. 2016; Reed et al. 2021; Templeton et al. 2016). On the other hand, assuming the simulated intruder is located, noise may interfere with the signaling behaviors of subjects which may induce them to resort to higher physical aggression such as closer approach (e.g., Grabarcyzk and Gill 2019). Thus, the differences in the findings may be due in part to differences in experimental designs, particularly with respect to the presentation of the noise stimulus (e.g., type of noise, location of noise relative to the conspecific stimulus etc.).

The experimental noise in our study represents a transient increase in noise to a high amplitude that coincides with the need to confront a territorial intruder. Thus, the effects we see are responses to acute noises, while urban birds would experience varying noise levels due to cars passing as well as daily patterns of human activity (Gill et al. 2017). Our study was explicitly designed to study plastic responses to acute increases in noise and therefore the results may not apply to chronic but varying amounts of noise. Nevertheless, the situation we simulated is a realistic one: urban wildlife has to deal with transient increases in noise such as this regularly (e.g. when a park worker uses a leafblower to clean trails or when a lawnmower works nearby).

### Change in multi-modal signals with noise

We found that European robins changed their signaling behaviors in response to noise. In the acoustic modality, urban but not rural robins decreased their song rates, while birds in neither habitat changed their song length. From the perspective of the multi-modal shift hypothesis, a decrease in signals in auditory modality was expected to coincide with an increase in visual signals. We did not find this second effect. Two caveats are worth mentioning here: First, we were not able to carry out the experiment blindly with respect to habitat or noise treatment, which may have biased our observations of visual displays. Clearly, we could not blind observers regarding which type of habitat they were in, and “blinding” observers with respect to noise treatment (e.g., by using noise-canceling headphones) would have made keeping track of the vocal behaviors almost impossible. In any case, given the pattern of results that tend to the opposite direction of our expectations however, we believe observer bias is unlikely to be an issue here.

Second, we could not examine individual level plasticity in visual signaling, because of a significant order effect in visual signals: visual displays were mostly used in the first trials only. We do not have a good hypothesis as to why we found such an order effect. It is possible that our stationary 3D model ceased to be a good visual stimulus by the time of the second trial, leading to a decrease in visual signaling. It is also possible that the second trials may have represented a lesser threat to the territory owner (given that it simulates the return of a previously retreating individual) thus eliciting lower threat signals. Lack (1939) noted that repeated presentations of a taxidermic mount quickly leads the lowered responses which he interpreted as the lack of realistic response of the immobile taxidermic mount. While something similar may be happening in our experiment, we note that that there was no order effect in other aggressive or vocal behaviors.

Whatever the reason, the order effect meant that we could only analyze the visual signals in the first trials for each subject, halving our sample size and precluding a within-subject comparison. The between individual comparison of visual signals among urban birds in the first trials yielded evidence in the opposite direction of what was expected: there was more visual signaling in trials without experimental noise than with experimental noise, although the difference was not significant, likely due to the small sample size. Nevertheless, this finding suggests that while urban birds decrease their acoustic signaling effort, they do not necessarily depend on visual signals as a back-up as predicted by the multi-modal shift hypothesis.

Our finding is similar to that of a study on European robins by Zwart et al (2016) which reported slightly lower rates of visual threat signals with experimental noise than without noise. Another study on song sparrows found no effect of experimental noise during territorial defense on overall rates of wing waves, a visual threat signal (Akçay and Beecher, 2019), even though urban song sparrows show higher rates of wing waves than rural birds when controlling for the total number of acoustic and visual threat signals (Akçay et al. 2020). These studies suggest that if urban noise causes a multi-modal shift in visual threat signals, it is unlikely to be due to immediate phenotypic plasticity.

Note that the fact that urban robins used visual displays more frequently than rural robins is consistent with a multimodal shift due to noise. This finding, however, is also consistent with the hypothesis that urban birds are simply more aggressive and therefore use visual threat signals more than rural birds (cf. Akçay et al. 2020). A valid comparison of the use of visual signals between urban and rural birds would need to somehow correct for aggressiveness. Thus, currently the evidence for a multi-modal shift in this species is relatively weak.

### Behavioral plasticity of urban vs. rural birds in response to noise

We had expected that urban birds would show a higher level of plasticity in their responses to experimentally presented noise compared to rural birds. This prediction was based on studies that showed prior experience with noise (as urban birds would have) leads to more directional plasticity in acoustic parameters of their song, by e.g. increasing the minimum frequency of their song in noise (Gentry et al. 2017; LaZerte et al. 2016). Instead, we found contrasting patterns of response to noise depending on the behavior measured: while in aggression scores, urban birds showed lower directional plasticity than rural birds (specifically, rural birds increased their approach distance in noise, while urban birds showed no difference), the opposite was true in song rates (urban birds decreased song rates in noise while rural birds did not show a difference between treatments).

These findings suggest that if there is a role of learning in determining responses, it may take different forms depending on the behavior. It is possible for instance that urban birds in general have learned to “sit out” transient increases in noise (such as the situation we simulated here) by reducing song rates and not increasing their approach towards opponents and visual signaling. Such a strategy may be adaptive, given that closely approaching an opponent and increasing visual threat signals likely would escalate an aggressive interaction to a fight (Searcy et al. 2006). In contrast, rural birds may show less plasticity in signing in noise and instead approach the opponent more closely during noise, because they haven’t had the opportunity to learn how to deal with transient increases in noise. Thus, the contrasting patterns found in urban and rural birds may still indicate the role of prior experience with noise, although clearly more controlled experiments are needed to further test this hypothesis.

### Conclusion

In summary, our results showed important differences in how urban and rural robins respond to noise during aggressive interactions. These results suggest that the ambient noise levels experienced by animals is an important factor in determining their responses to transient increases in noise.

## Supporting information

Supplementary Materials (with code)

## Disclosure of potential conflicts of interest

The authors declare that they have no conflicts of interest.

Research involving Human Participants and/or Animals

All procedures used in this study follow the ASAB/ABS guidelines for the treatment of animals in behavioral research and teaching. Subjects were not captured or handled before, during or after any of the trials. Time spent within a territory did not exceed 15 minutes per trial, and 30 minutes per day.

## Informed consent

The research did not involve human participants.

## Data availability statement

The raw data and the R-code to reproduce the analyses reported in the manuscript is available as a supplementary material and will be deposited at a public repository before the manuscript is published.

## Acknowledgements

This study was funded by the British Ornithological Union Small Ornithological Research Grant to çÖ.

