## Supplementary Materials for "Aggression and multimodal signaling in noise in a common urban songbird"

### Onsaletal2022_Analysis

CaglaOnsal

4/19/2022

#### Noise Playback Power Spectrum (Figure S1)

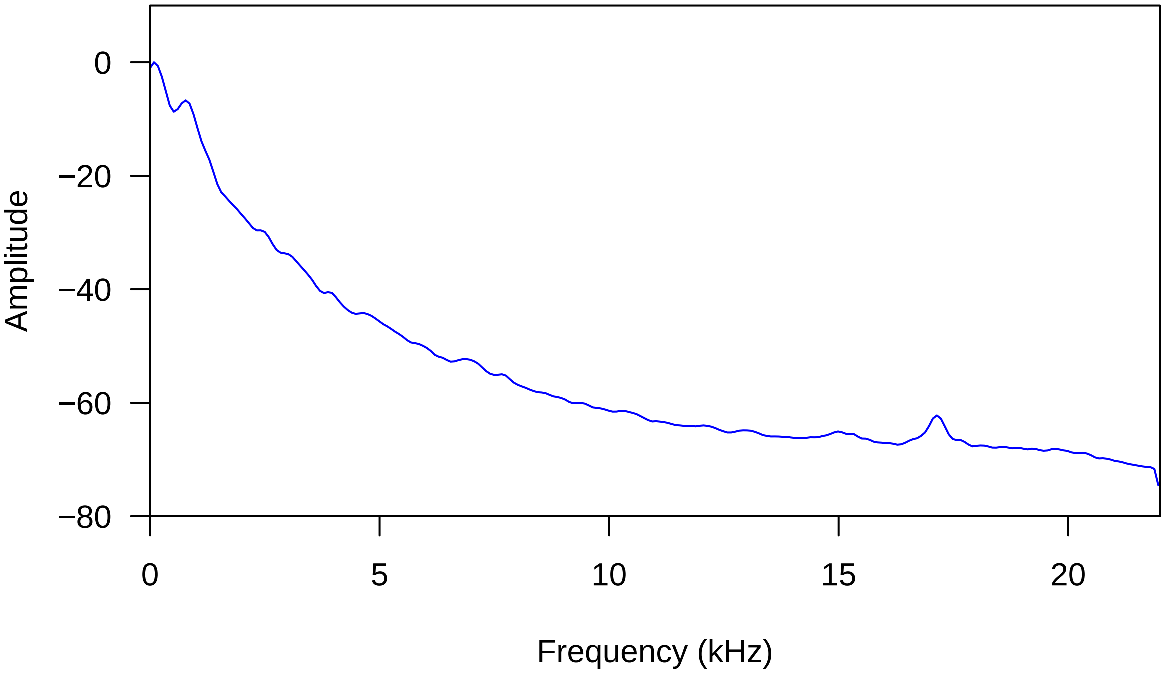

#### Song Spectrogram & Power spectrum (Figure S2 and S3)

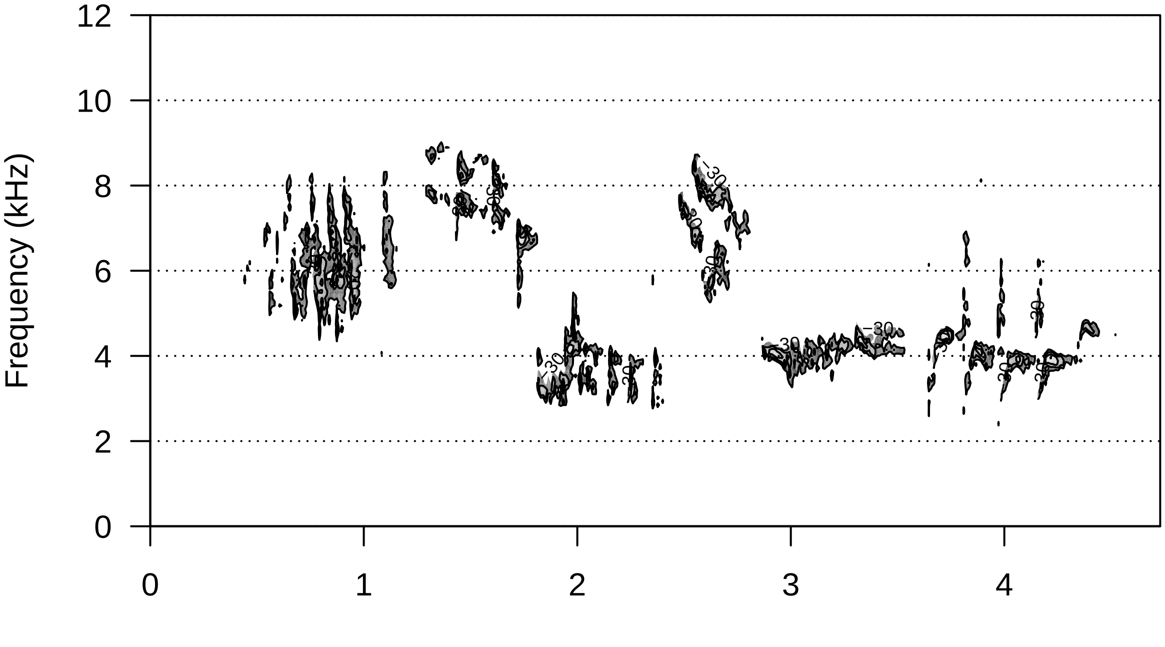


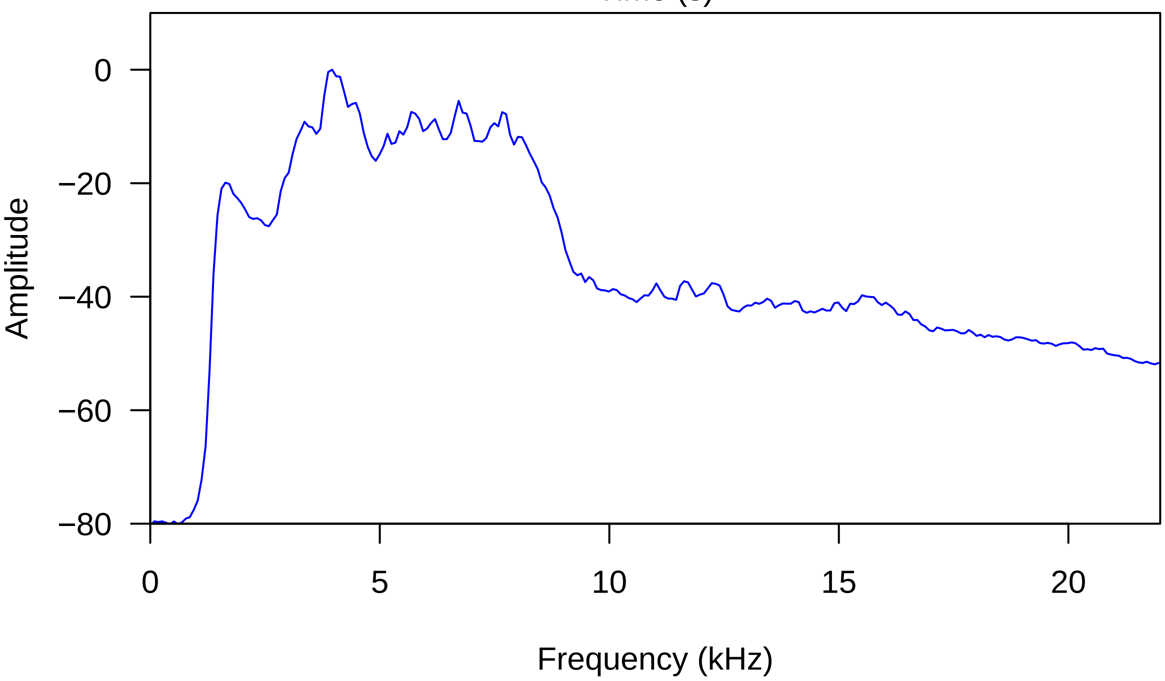


#### Principal Component Analyses on spatial variables

##### Correlations

###### Flight rate and proportion of time spent within 1m

```
##
##  Pearson's product-moment correlation
##
```

```
## data:  Robin2021$FlightRate and Robin2021$prop1m
## t = 2.2194, df = 40, p-value = 0.03219
## alternative hypothesis: true correlation is not equal to 0
## 95 percent confidence interval:
##  0.03023823 0.57698971
## sample estimates:
##       cor
## 0.3311264
```

**Flight rate and closest approach(m)**

```
##
##  Pearson's product-moment correlation
##
## data:  Robin2021$FlightRate and Robin2021$ClosestApproachM
## t = -2.3596, df = 40, p-value = 0.02326
## alternative hypothesis: true correlation is not equal to 0
## 95 percent confidence interval:
##  -0.59072113 -0.05103777
## sample estimates:
##       cor
## -0.349547
```

**Closest approach(m) and proportion of time spent within 1m**

```
##
##  Pearson's product-moment correlation
##
## data:  Robin2021$ClosestApproachM and Robin2021$prop5m
## t = -4.5283, df = 40, p-value = 5.25e-05
## alternative hypothesis: true correlation is not equal to 0
## 95 percent confidence interval:
##  -0.7528763 -0.3380341
## sample estimates:
##        cor
## -0.5821553
```

#### Aggression scores (PCA1) distribution (Figure S4)

```
## Principal Components Analysis
## Call: principal(r = PhysicalMeasures, nfactors = 1, residuals = FALSE,
##     rotate = "none", covar = FALSE)
## Standardized loadings (pattern matrix) based upon correlation matrix
##                             PC1   h2   u2 com
## Robin2021$FlightRate        0.65 0.42 0.58   1
## Robin2021$ClosestApproachM -0.85 0.73 0.27   1
## Robin2021$prop5m            0.83 0.68 0.32   1
##
##                  PC1
## SS loadings    1.83
## Proportion Var 0.61
##
## Mean item complexity =  1
## Test of the hypothesis that 1 component is sufficient.
##
```

```
## The root mean square of the residuals (RMSR) is  0.2
##  with the empirical chi square  9.75  with prob <  NA
##
## Fit based upon off diagonal values = 0.79
```

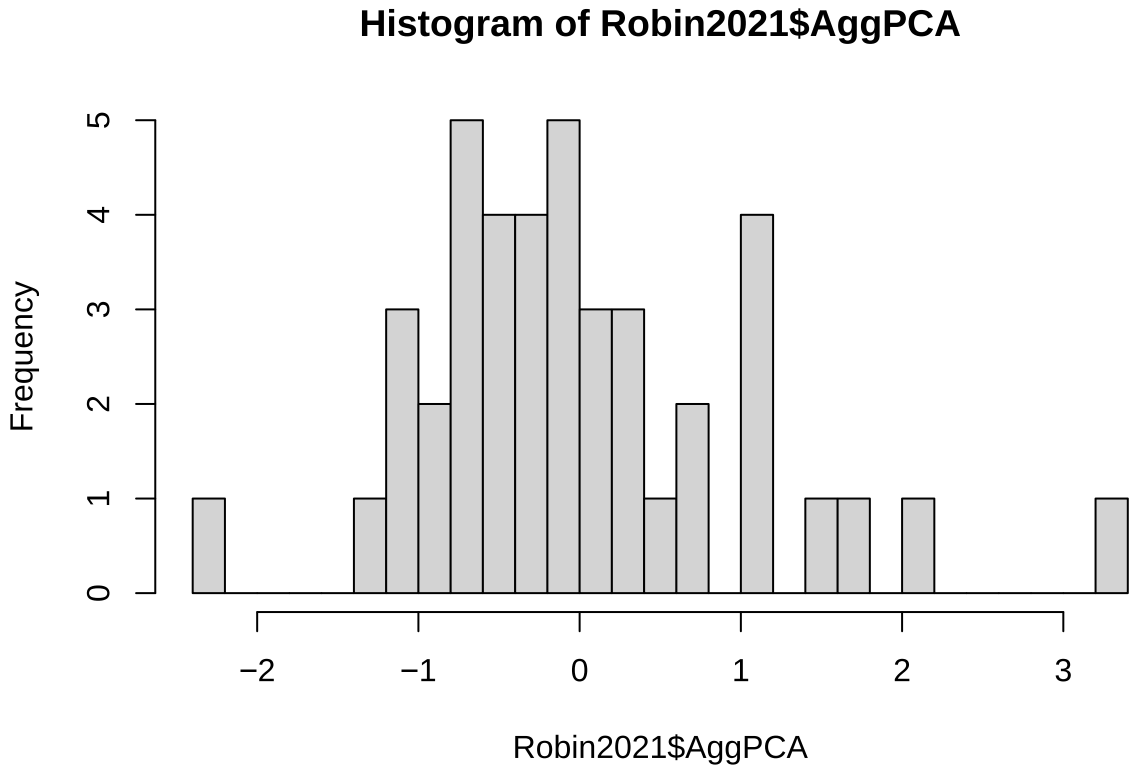


#### Ambient Noise

##### Ambient Noise and Habitat, LMM

```
## Warning: NAs introduced by coercion
```

```
## Linear mixed-effects model fit by REML
##   Data: Robin2021
##        AIC      BIC    logLik
##   204.1462 210.5899 -98.07312
##
## Random effects:
##  Formula: ~1 | MaleID
##         (Intercept) Residual
## StdDev:     5.27673 1.324208
##
## Fixed effects:  Average ~ Habitat
##                  Value Std.Error DF   t-value p-value
## (Intercept)  39.69781  1.550891 19 25.596782  0.0000
## HabitatUrban  8.96880  2.367907 19  3.787648  0.0012
##  Correlation:
##              (Intr)
## HabitatUrban -0.655
##
## Standardized Within-Group Residuals:
```

```
##         Min          Q1         Med          Q3         Max
## -1.31435938 -0.58807449 -0.05934325  0.59745834  1.62135675
##
## Number of Observations: 39
## Number of Groups: 21
```

```
##             numDF denDF   F-value p-value
## (Intercept)     1    19 1380.5883  <.0001
## Habitat         1    19   14.3463  0.0012
```

##### Repeatability of noise measurements

```
## Warning in if (datatype == "Gaussian") {: the condition has length > 1 and only
## the first element will be used
```

```
## Warning in rptGaussian(formula, grname, data, CI, nboot, npermut, parallel, : 3
## rows containing missing values were removed
```

```
## Bootstrap Progress:
```

```
##
##
## Repeatability estimation using the lmm method
##
## Repeatability for MaleID
## R  = 0.964
## SE = 0.019
## CI = [0.912, 0.985]
## P  = 2.56e-12 [LRT]
##      NA [Permutation]
```

##### Ambient noise by habitat, Mean and SD

###### Urban

```
## [1] 48.98088
```

```
## [1] 7.071437
```

###### Rural

```
## [1] 39.94472
```

```
## [1] 3.570615
```

#### Order effect

##### Order Effect on song rates

```
## Linear mixed-effects model fit by REML
##   Data: Robin2021
##        AIC      BIC    logLik
##   188.9021 195.6576 -90.45104
##
## Random effects:
##  Formula: ~1 | MaleID
##         (Intercept) Residual
## StdDev:    0.898853 1.972697
```

```
##
## Fixed effects:  SongRateTrial ~ Order
##                  Value Std.Error DF   t-value p-value
## (Intercept)  8.396825 0.4730585 20 17.750079  0.0000
## Ordersecond -0.682540 0.6087876 20 -1.121146  0.2755
##  Correlation:
##             (Intr)
## Ordersecond -0.643
##
## Standardized Within-Group Residuals:
##        Min         Q1        Med         Q3        Max
## -2.4872641 -0.4720833  0.2082500  0.6657558  2.2505976
##
## Number of Observations: 42
## Number of Groups: 21
```

```
## Analysis of Deviance Table (Type III tests)
##
## Response: SongRateTrial
##               Chisq Df Pr(>Chisq)
## (Intercept) 315.065  1     <2e-16 ***
## Order         1.257  1     0.2622
## ---
## Signif. codes:  0 '***' 0.001 '**' 0.01 '*' 0.05 '.' 0.1 ' ' 1
```

##### Order effect on aggression scores (PCA1)

```
## Linear mixed-effects model fit by REML
##   Data: Robin2021
##        AIC      BIC    logLik
##   110.5753 117.3308 -51.28765
##
## Random effects:
##  Formula: ~1 | MaleID
##         (Intercept)  Residual
## StdDev:   0.8792328 0.4889573
##
## Fixed effects:  AggPCA ~ Order
##                   Value Std.Error DF    t-value p-value
## (Intercept)  0.1107159 0.2195373 20  0.5043147  0.6196
## Ordersecond -0.2214318 0.1508955 20 -1.4674515  0.1578
##  Correlation:
##             (Intr)
## Ordersecond -0.344
##
## Standardized Within-Group Residuals:
##         Min          Q1         Med          Q3         Max
## -1.64117543 -0.40591283  0.01055451  0.46065047  1.71283940
##
## Number of Observations: 42
## Number of Groups: 21
```

```
## Analysis of Deviance Table (Type III tests)
##
## Response: AggPCA
```

```
##              Chisq Df Pr(>Chisq)
## (Intercept) 0.2543  1     0.6140
## Order       2.1534  1     0.1423
```

##### Order effect on visual displays

```
## Generalized linear mixed model fit by maximum likelihood (Laplace
##   Approximation) [glmerMod]
##  Family: binomial  ( logit )
## Formula: VisualDisplay ~ Order + (1 | MaleID)
##    Data: Robin2021
##
##      AIC      BIC   logLik deviance df.resid
##     31.9     37.1    -12.9     25.9       39
##
## Scaled residuals:
##       Min        1Q    Median        3Q       Max
## -0.006653 -0.003376 -0.000002 -0.000002  0.096861
##
## Random effects:
##  Groups Name        Variance Std.Dev.
##  MaleID (Intercept) 3371     58.06
## Number of obs: 42, groups:  MaleID, 21
##
## Fixed effects:
##             Estimate Std. Error z value Pr(>|z|)
## (Intercept)  -11.344      3.358  -3.379 0.000729 ***
## Ordersecond  -15.323      6.221  -2.463 0.013778 *
## ---
## Signif. codes:  0 '***' 0.001 '**' 0.01 '*' 0.05 '.' 0.1 ' ' 1
##
## Correlation of Fixed Effects:
##             (Intr)
## Ordersecond 0.474
```

```
## Analysis of Deviance Table (Type III Wald chisquare tests)
##
## Response: VisualDisplay
##               Chisq Df Pr(>Chisq)
## (Intercept) 11.4147  1  0.0007286 ***
## Order        6.0664  1  0.0137778 *
## ---
## Signif. codes:  0 '***' 0.001 '**' 0.01 '*' 0.05 '.' 0.1 ' ' 1
```

#### Aggression score analyses

##### Linear mixed models on Aggression scores, with habitat and treatment as predictors (Table 3)

```
## Linear mixed-effects model fit by REML
##   Data: Robin2021
##        AIC      BIC    logLik
##   105.9003 115.7259 -46.95017
##
```

```
## Random effects:
##  Formula: ~1 | MaleID
##         (Intercept)  Residual
## StdDev:   0.7549836 0.4650489
##
## Fixed effects:  AggPCA ~ Habitat * Condition
##                                  Value Std.Error DF   t-value p-value
## (Intercept)                 -0.5826742 0.2559737 19 -2.276304  0.0346
## HabitatUrban                 1.3208684 0.3910064 19  3.378125  0.0032
## Conditionnoise               0.3229697 0.1898554 19  1.701135  0.1052
## HabitatUrban:Conditionnoise -0.6761866 0.2900089 19 -2.331606  0.0309
##  Correlation:
##                             (Intr) HbttUr Cndtnn
## HabitatUrban                -0.655
## Conditionnoise              -0.371  0.243
## HabitatUrban:Conditionnoise  0.243 -0.371 -0.655
##
## Standardized Within-Group Residuals:
##        Min         Q1        Med         Q3        Max
## -1.6836177 -0.3376003  0.0175153  0.3696283  1.9579139
##
## Number of Observations: 42
## Number of Groups: 21
```

```
## Analysis of Deviance Table (Type III tests)
##
## Response: AggPCA
##                     Chisq Df Pr(>Chisq)
## (Intercept)        5.1816  1  0.0228278 *
## Habitat           11.4117  1  0.0007298 ***
## Condition          2.8939  1  0.0889176 .
## Habitat:Condition  5.4364  1  0.0197214 *
## ---
## Signif. codes:  0 '***' 0.001 '**' 0.01 '*' 0.05 '.' 0.1 ' ' 1
```

**LMM on aggression scores with noise treatment as predictor, Rural Subset**

```
## Linear mixed-effects model fit by REML
##   Data: Robin2021
##   Subset: Habitat == "Rural"
##        AIC      BIC    logLik
##   50.36621 54.73038 -21.18311
##
## Random effects:
##  Formula: ~1 | MaleID
##         (Intercept)  Residual
## StdDev:   0.6600083 0.3242974
##
## Fixed effects:  AggPCA ~ Condition
##                     Value Std.Error DF   t-value p-value
## (Intercept)    -0.5826742 0.2122851 11 -2.744772  0.0191
## Conditionnoise  0.3229697 0.1323939 11  2.439462  0.0329
##  Correlation:
##                (Intr)
## Conditionnoise -0.312
```

```
##
## Standardized Within-Group Residuals:
##         Min          Q1         Med          Q3         Max
## -2.05344186 -0.32001375  0.08331217  0.26491375  1.30934217
##
## Number of Observations: 24
## Number of Groups: 12
```

**LMM on aggression scores with noise treatment as predictor, Urban Subset**

```
## Linear mixed-effects model fit by REML
##   Data: Robin2021
##   Subset: Habitat == "Urban"
##        AIC      BIC    logLik
##   54.86982 57.96017 -23.43491
##
## Random effects:
##  Formula: ~1 | MaleID
##         (Intercept)  Residual
## StdDev:   0.8687837 0.6074828
##
## Fixed effects:  AggPCA ~ Condition
##                      Value Std.Error DF   t-value p-value
## (Intercept)      0.7381942 0.3533680  8  2.089024  0.0701
## Conditionnoise -0.3532169 0.2863701  8 -1.233428  0.2524
##  Correlation:
##                (Intr)
## Conditionnoise -0.405
##
## Standardized Within-Group Residuals:
##         Min          Q1         Med          Q3         Max
## -1.26451755 -0.53660150 -0.02836087  0.46799618  1.52319947
##
## Number of Observations: 18
## Number of Groups: 9
```

**T test, aggression scores in urban and rural birds in noise treatment**

```
##
##  Welch Two Sample t-test
##
## data:  AggPCA by Habitat
## t = -1.8348, df = 12.361, p-value = 0.04536
## alternative hypothesis: true difference in means between group Rural and group Urban is less than 0
## 95 percent confidence interval:
##         -Inf -0.01995465
## sample estimates:
## mean in group Rural mean in group Urban
##          -0.2597045           0.3849773
```

**Models with Individual Spatial Measures**

**Closest Approach Distance (m)**

```
## Linear mixed-effects model fit by REML
```

```
##   Data: Robin2021
##       AIC      BIC    logLik
##   230.135 239.9605 -109.0675
##
## Random effects:
##  Formula: ~1 | MaleID
##         (Intercept) Residual
## StdDev:    3.392613 2.607776
##
## Fixed effects:  ClosestApproachM ~ Habitat * Condition
##                                Value Std.Error DF   t-value p-value
## (Intercept)                10.750000  1.235257 19  8.702642  0.0000
## HabitatUrban               -6.616667  1.886886 19 -3.506659  0.0024
## Conditionnoise             -3.583333  1.064620 19 -3.365833  0.0032
## HabitatUrban:Conditionnoise  3.827778  1.626234 19  2.353768  0.0295
##  Correlation:
##                            (Intr) HbttUr Cndtnn
## HabitatUrban               -0.655
## Conditionnoise             -0.431  0.282
## HabitatUrban:Conditionnoise  0.282 -0.431 -0.655
##
## Standardized Within-Group Residuals:
##         Min          Q1         Med          Q3         Max
## -1.44199595 -0.38285212 -0.03918232  0.39303293  2.93593593
##
## Number of Observations: 42
## Number of Groups: 21
```

```
## Analysis of Deviance Table (Type III tests)
##
## Response: ClosestApproachM
##                     Chisq Df Pr(>Chisq)
## (Intercept)       75.7360  1  < 2.2e-16 ***
## Habitat           12.2967  1  0.0004538 ***
## Condition         11.3288  1  0.0007631 ***
## Habitat:Condition  5.5402  1  0.0185842 *
## ---
## Signif. codes:  0 '***' 0.001 '**' 0.01 '*' 0.05 '.' 0.1 ' ' 1
```

**Proportion of time spent within 5m of the model**

```
## Linear mixed-effects model fit by REML
##   Data: Robin2021
##        AIC      BIC    logLik
##   38.72572 48.55123 -13.36286
##
## Random effects:
##  Formula: ~1 | MaleID
##         (Intercept)  Residual
## StdDev:   0.2194376 0.2368401
##
## Fixed effects:  prop5m ~ Habitat * Condition
##                                 Value  Std.Error DF    t-value p-value
## (Intercept)                 0.1129630 0.09320501 19  1.2119838  0.2404
## HabitatUrban                0.3660494 0.14237301 19  2.5710588  0.0187
```

```
## Conditionnoise              -0.0509259 0.09668957 19 -0.5266951  0.6045
## HabitatUrban:Conditionnoise -0.0854938 0.14769576 19 -0.5788509  0.5695
##  Correlation:
##                             (Intr) HbttUr Cndtnn
## HabitatUrban                -0.655
## Conditionnoise              -0.519  0.340
## HabitatUrban:Conditionnoise  0.340 -0.519 -0.655
##
## Standardized Within-Group Residuals:
##         Min          Q1         Med          Q3         Max
## -1.48439572 -0.29341059 -0.08769457  0.12624752  2.14809796
##
## Number of Observations: 42
## Number of Groups: 21
```

```
## Analysis of Deviance Table (Type III tests)
##
## Response: prop5m
##                   Chisq Df Pr(>Chisq)
## (Intercept)       1.4689  1    0.22552
## Habitat           6.6103  1    0.01014 *
## Condition         0.2774  1    0.59841
## Habitat:Condition 0.3351  1    0.56269
## ---
## Signif. codes:  0 '***' 0.001 '**' 0.01 '*' 0.05 '.' 0.1 ' ' 1
```

##### Flight Rate

```
## Linear mixed-effects model fit by REML
##   Data: Robin2021
##        AIC      BIC    logLik
##   168.7912 178.6168 -78.39562
##
## Random effects:
##  Formula: ~1 | MaleID
##         (Intercept) Residual
## StdDev:      1.1967 1.320389
##
## Fixed effects:  FlightRate ~ Habitat * Condition
##                                 Value Std.Error DF   t-value p-value
## (Intercept)                  1.222222 0.5144184 19  2.375930  0.0282
## HabitatUrban                 1.148148 0.7857871 19  1.461144  0.1603
## Conditionnoise               0.250000 0.5390464 19  0.463782  0.6481
## HabitatUrban:Conditionnoise -1.027778 0.8234070 19 -1.248201  0.2271
##  Correlation:
##                             (Intr) HbttUr Cndtnn
## HabitatUrban                -0.655
## Conditionnoise              -0.524  0.343
## HabitatUrban:Conditionnoise  0.343 -0.524 -0.655
##
## Standardized Within-Group Residuals:
##        Min         Q1        Med         Q3        Max
## -0.9975790 -0.4532515 -0.1859754  0.3122705  3.2416885
##
## Number of Observations: 42
```

```
## Number of Groups: 21
```

```
## Analysis of Deviance Table (Type III tests)
##
## Response: FlightRate
##                    Chisq Df Pr(>Chisq)
## (Intercept)       5.6450  1     0.0175 *
## Habitat           2.1349  1     0.1440
## Condition         0.2151  1     0.6428
## Habitat:Condition 1.5580  1     0.2120
## ---
## Signif. codes:  0 '***' 0.001 '**' 0.01 '*' 0.05 '.' 0.1 ' ' 1
```

#Song rate ## LMM on song rates, with habitat and treatment as predictor variables (Table 3)

```
## Linear mixed-effects model fit by REML
##   Data: Robin2021
##        AIC      BIC    logLik
##   180.6677 190.4932 -84.33385
##
## Random effects:
##  Formula: ~1 | MaleID
##         (Intercept) Residual
## StdDev:    1.260692 1.611867
##
## Fixed effects:  SongRateTrial ~ Habitat * Condition
##                                 Value Std.Error DF   t-value p-value
## (Intercept)                  8.444444 0.5907241 19 14.295074  0.0000
## HabitatUrban                 0.814815 0.9023460 19  0.902996  0.3778
## Conditionnoise              -0.611111 0.6580420 19 -0.928681  0.3647
## HabitatUrban:Conditionnoise -2.018519 1.0051757 19 -2.008125  0.0591
##  Correlation:
##                             (Intr) HbttUr Cndtnn
## HabitatUrban                -0.655
## Conditionnoise              -0.557  0.365
## HabitatUrban:Conditionnoise  0.365 -0.557 -0.655
##
## Standardized Within-Group Residuals:
##         Min          Q1         Med          Q3         Max
## -2.46977565 -0.44814419  0.02979613  0.44142364  1.66060285
##
## Number of Observations: 42
## Number of Groups: 21
```

```
## Analysis of Deviance Table (Type III tests)
##
## Response: SongRateTrial
##                      Chisq Df Pr(>Chisq)
## (Intercept)       204.3491  1    < 2e-16 ***
## Habitat             0.8154  1    0.36653
## Condition           0.8624  1    0.35305
## Habitat:Condition   4.0326  1    0.04463 *
## ---
## Signif. codes:  0 '***' 0.001 '**' 0.01 '*' 0.05 '.' 0.1 ' ' 1
```

**LMM on song rates, with noise treatment as predictor variable, Urban subset**

```
## Linear mixed-effects model fit by REML
##   Data: Robin2021
##   Subset: Habitat == "Urban"
##        AIC     BIC    logLik
##   77.11524 80.2056 -34.55762
##
## Random effects:
##  Formula: ~1 | MaleID
##         (Intercept) Residual
## StdDev:   0.4530418 1.773432
##
## Fixed effects:  SongRateTrial ~ Condition
##                    Value Std.Error DF   t-value p-value
## (Intercept)     9.259259 0.6101283  8 15.175922  0.0000
## Conditionnoise -2.629630 0.8360040  8 -3.145475  0.0137
##  Correlation:
##                (Intr)
## Conditionnoise -0.685
##
## Standardized Within-Group Residuals:
##         Min          Q1         Med          Q3         Max
## -1.52328188 -0.48964768 -0.05582765  0.31825570  1.95462053
##
## Number of Observations: 18
## Number of Groups: 9
```

```
## Analysis of Deviance Table (Type III tests)
##
## Response: SongRateTrial
##               Chisq Df Pr(>Chisq)
## (Intercept) 230.309  1  < 2.2e-16 ***
## Condition     9.894  1   0.001658 **
## ---
## Signif. codes:  0 '***' 0.001 '**' 0.01 '*' 0.05 '.' 0.1 ' ' 1
```

**LMM on song rates, with noise treatment as predictor variable, Rural Subset**

```
## Linear mixed-effects model fit by REML
##   Data: Robin2021
##   Subset: Habitat == "Rural"
##        AIC      BIC    logLik
##   106.0826 110.4468 -49.04129
##
## Random effects:
##  Formula: ~1 | MaleID
##         (Intercept) Residual
## StdDev:    1.611198 1.483353
##
## Fixed effects:  SongRateTrial ~ Condition
##                    Value Std.Error DF  t-value p-value
## (Intercept)     8.444444 0.6322115 11 13.35699  0.0000
## Conditionnoise -0.611111 0.6055764 11 -1.00914  0.3346
##  Correlation:
```

```
##                (Intr)
## Conditionnoise -0.479
##
## Standardized Within-Group Residuals:
##        Min         Q1        Med         Q3        Max
## -2.2764595 -0.2585715  0.0236137  0.5459746  0.9311823
##
## Number of Observations: 24
## Number of Groups: 12
```

```
## Analysis of Deviance Table (Type III tests)
##
## Response: SongRateTrial
##                Chisq Df Pr(>Chisq)
## (Intercept) 178.4093  1     <2e-16 ***
## Condition     1.0184  1     0.3129
## ---
## Signif. codes:  0 '***' 0.001 '**' 0.01 '*' 0.05 '.' 0.1 ' ' 1
```

#Visual Signals

#### GLM on Visual Signals in the first trials, habitat and treatment as predictor variables

```
##
## Call:
## glm(formula = VisualDisplay ~ Habitat * Condition, family = binomial,
##     data = Robin2021, subset = Order == "first")
##
## Deviance Residuals:
##      Min        1Q    Median        3Q       Max
## -0.75853  -0.60386  -0.60386   0.00013   1.89302
##
## Coefficients:
##                              Estimate Std. Error z value Pr(>|z|)
## (Intercept)                -1.609e+00  1.095e+00  -1.469    0.142
## HabitatUrban                2.018e+01  2.917e+03   0.007    0.994
## Conditionnoise              1.435e-15  1.549e+00   0.000    1.000
## HabitatUrban:Conditionnoise -1.966e+01  2.917e+03  -0.007    0.995
##
## (Dispersion parameter for binomial family taken to be 1)
##
##     Null deviance: 27.910  on 20  degrees of freedom
## Residual deviance: 15.312  on 17  degrees of freedom
## AIC: 23.312
##
## Number of Fisher Scoring iterations: 17
```

```
## Analysis of Deviance Table (Type III tests)
##
## Response: VisualDisplay
##                   LR Chisq Df Pr(>Chisq)
## Habitat             9.7515  1   0.001792 **
## Condition           0.0000  1   1.000000
## Habitat:Condition   3.7288  1   0.053480 .
```

```
## ---
## Signif. codes:  0 '***' 0.001 '**' 0.01 '*' 0.05 '.' 0.1 ' ' 1
```

**GLM on Visual Signals in the first trials, treatment as predictor variable, Urban Subset**

```
##
## Call:
## glm(formula = VisualDisplay ~ Condition, family = binomial, data = Robin2021,
##     subset = Order == "first" & Habitat == "Urban")
##
## Deviance Residuals:
##      Min        1Q    Median        3Q       Max
## -0.75853  -0.75853   0.00005   0.00005   1.66511
##
## Coefficients:
##                Estimate Std. Error z value Pr(>|z|)
## (Intercept)       20.57    7929.26   0.003    0.998
## Conditionnoise   -21.66    7929.26  -0.003    0.998
##
## (Dispersion parameter for binomial family taken to be 1)
##
##     Null deviance: 11.4573  on 8  degrees of freedom
## Residual deviance:  4.4987  on 7  degrees of freedom
## AIC: 8.4987
##
## Number of Fisher Scoring iterations: 19
```

```
## Analysis of Deviance Table
##
## Model: binomial, link: logit
##
## Response: VisualDisplay
##
## Terms added sequentially (first to last)
##
##
##           Df Deviance Resid. Df Resid. Dev
## NULL                          8    11.4573
## Condition  1   6.9586         7     4.4987
```

#### GLM on whether aggression scores predict visual displays (binomial response variable)

```
##
## Call:
## glm(formula = VisualDisplay ~ AggPCA, family = binomial, data = Robin2021,
##     subset = Order == "first")
##
## Deviance Residuals:
##     Min       1Q   Median       3Q      Max
## -1.1975  -0.4202  -0.1430   0.2025   2.1478
##
## Coefficients:
##             Estimate Std. Error z value Pr(>|z|)
## (Intercept)  -0.9017     0.7532  -1.197   0.2313
```

```
## AggPCA        3.3270     1.5002   2.218   0.0266 *
## ---
## Signif. codes:  0 '***' 0.001 '**' 0.01 '*' 0.05 '.' 0.1 ' ' 1
##
## (Dispersion parameter for binomial family taken to be 1)
##
##     Null deviance: 27.910  on 20  degrees of freedom
## Residual deviance: 11.821  on 19  degrees of freedom
## AIC: 15.821
##
## Number of Fisher Scoring iterations: 6

## Warning in anova.glm(VisAgg, type = 3): the following arguments to 'anova.glm'
## are invalid and dropped: list(type = 3)

## Analysis of Deviance Table
##
## Model: binomial, link: logit
##
## Response: VisualDisplay
##
## Terms added sequentially (first to last)
##
##
##        Df Deviance Resid. Df Resid. Dev
## NULL                       20     27.910
## AggPCA  1   16.089        19     11.821
```

### Song Length

#### Song length, habitat and treatment

```
## Linear mixed-effects model fit by REML
##   Data: Robin2021
##        AIC      BIC   logLik
##   62.00559 71.83111 -25.0028
##
## Random effects:
##  Formula: ~1 | MaleID
##         (Intercept)  Residual
## StdDev:   0.4053826 0.2693497
##
## Fixed effects:  SongLengthAvg ~ Habitat * Condition
##                                 Value Std.Error DF   t-value p-value
## (Intercept)                 1.8611417 0.1405004 19 13.246524  0.0000
## HabitatUrban                0.0424111 0.2146179 19  0.197612  0.8454
## Conditionnoise              0.0298425 0.1099616 19  0.271390  0.7890
## HabitatUrban:Conditionnoise 0.1054574 0.1679690 19  0.627839  0.5376
##  Correlation:
##                             (Intr) HbttUr Cndtnn
## HabitatUrban                -0.655
## Conditionnoise              -0.391  0.256
## HabitatUrban:Conditionnoise  0.256 -0.391 -0.655
##
## Standardized Within-Group Residuals:
```

```
##        Min         Q1        Med         Q3        Max 
## -1.8093423 -0.5218645 -0.1315417  0.5450639  2.2035506 
## 
## Number of Observations: 42
## Number of Groups: 21
```

```
##                   numDF denDF  F-value p-value
## (Intercept)           1    19 384.6253  <.0001
## Habitat               1    19   0.2320  0.6355
## Condition             1    19   0.8149  0.3780
## Habitat:Condition     1    19   0.3942  0.5376
```

### Figures

**Figure S5- Visual displays in the first trials are correlated with aggression scores**

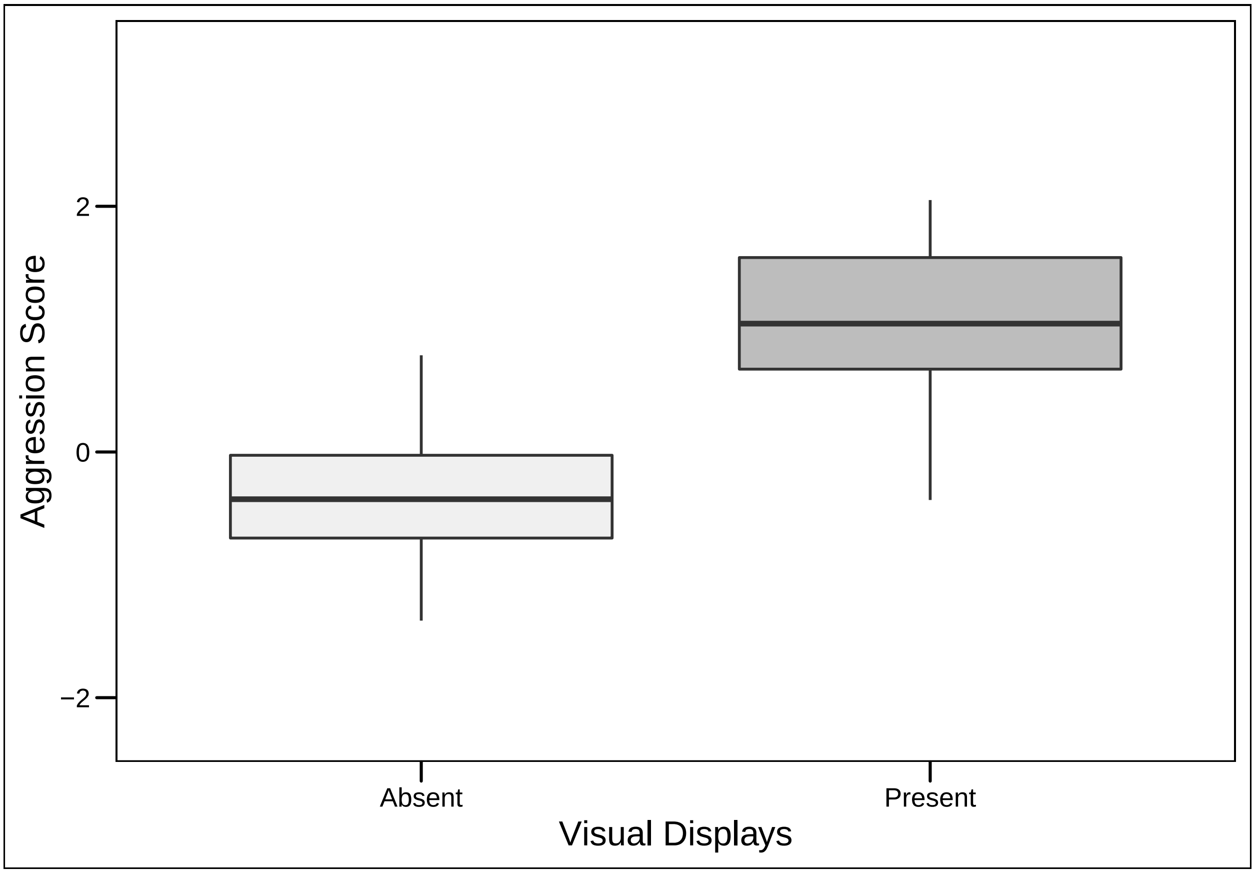

**Aggression, habitat and treatment (Figure 1a)**

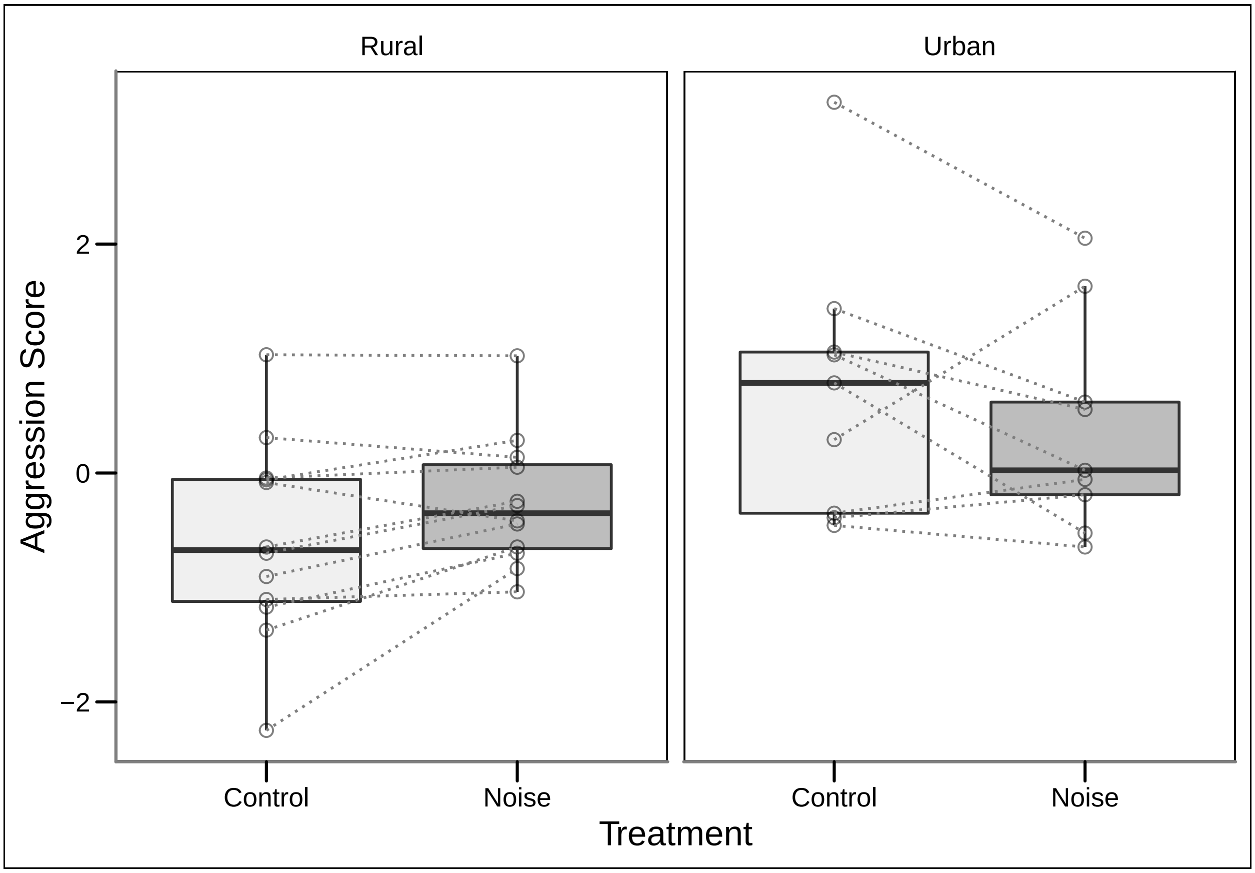

**Song rate, habitat and treatment (Figure 1b)**

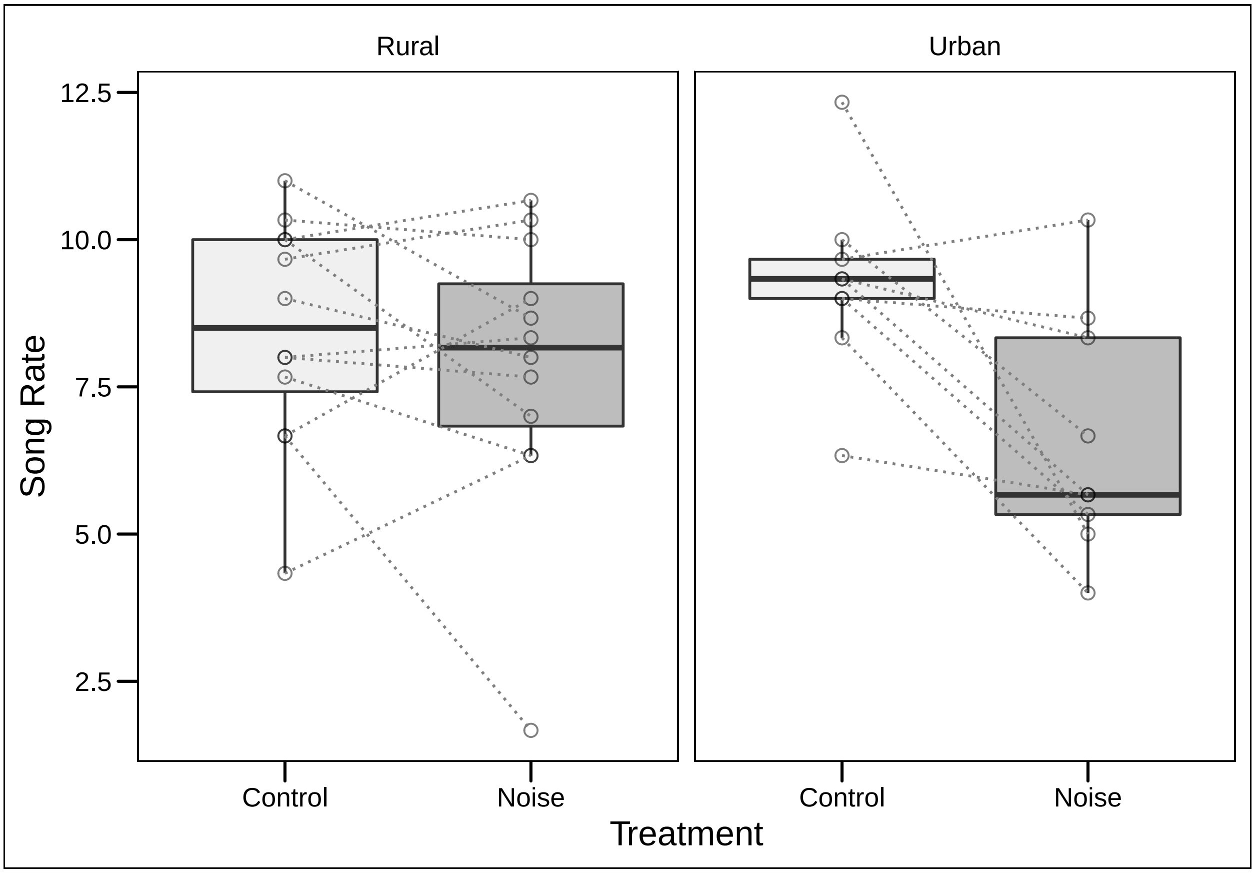

**Visual signals, habitat and treatment (Figure 2)**

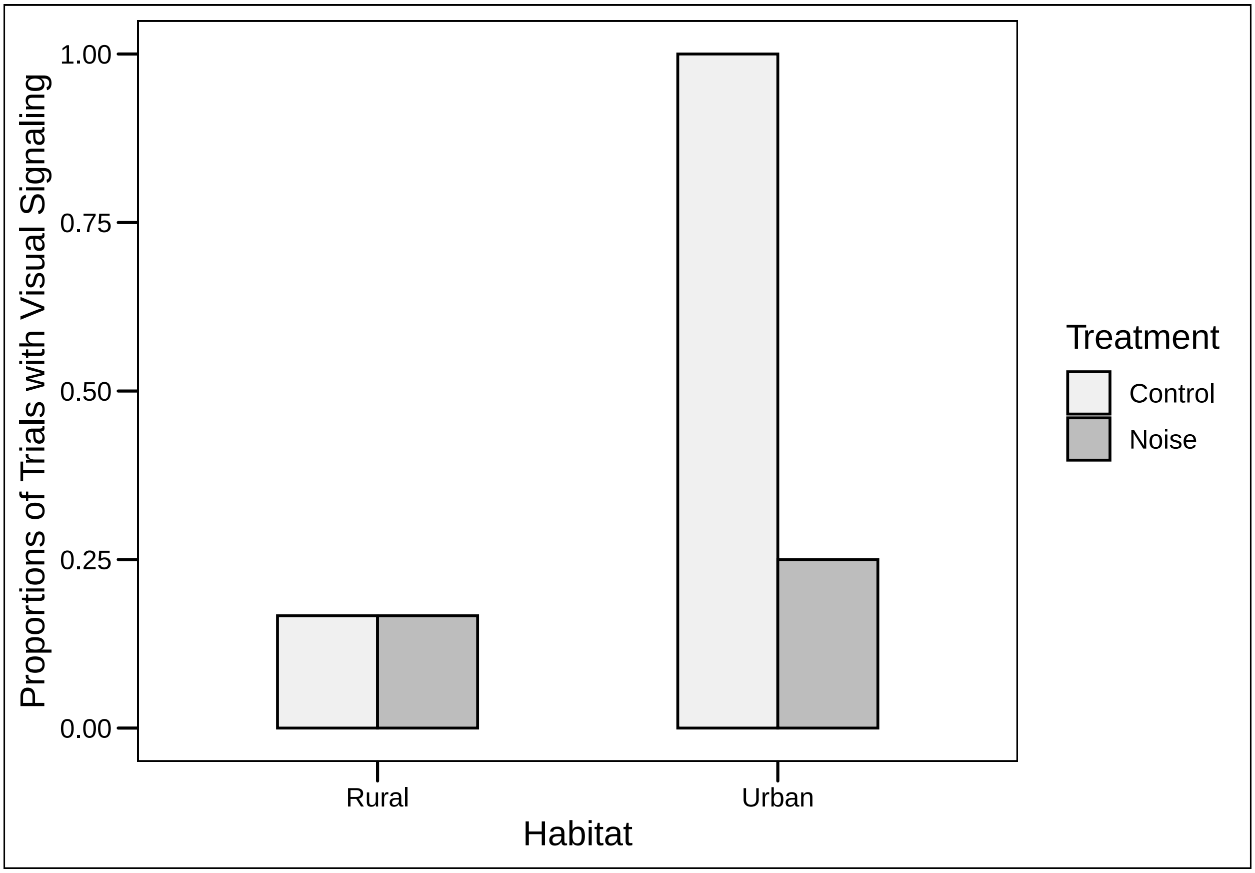

**Figure S6 - Visual Stimulus**

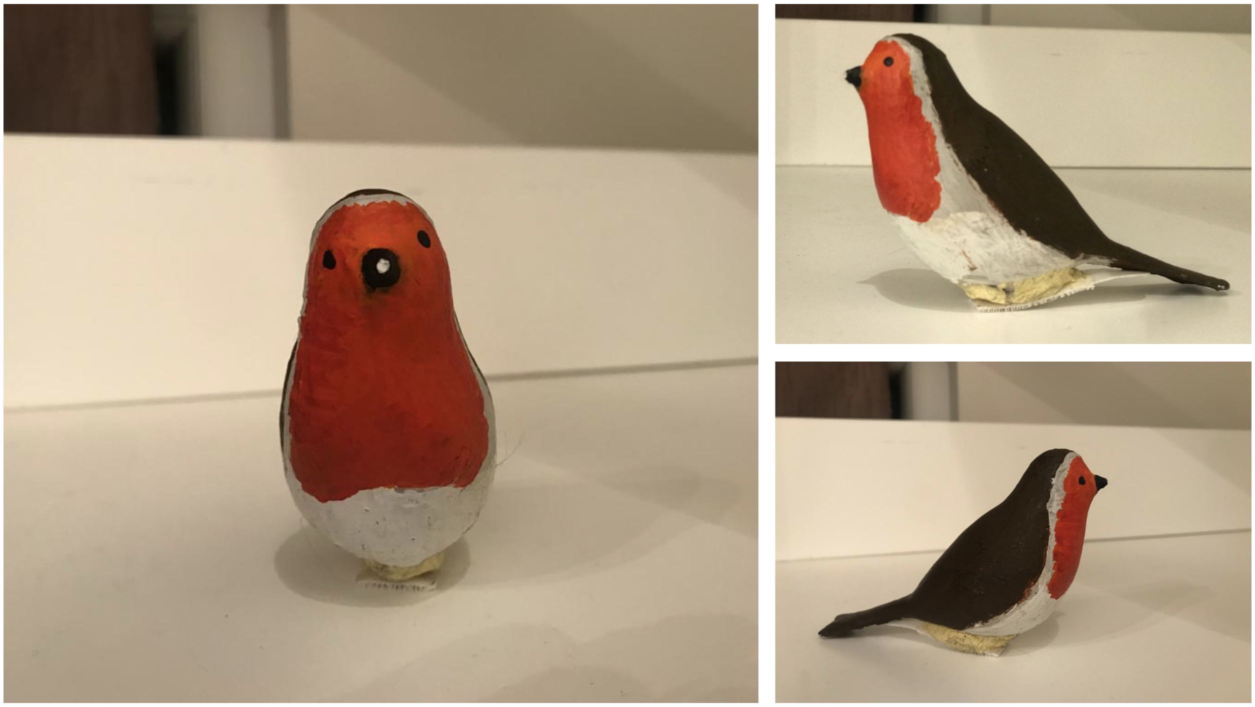

**S7 - A video collage of visual displays used by robins: https://youtu.be/CBuSxSc24Io**
