## Supplementary Materials (with code) for "Aggression and multimodal signaling in noise in a common urban songbird"

### Onsaletal2022_Code

CaglaOnsal

4/19/2022

### Aggression Score

### Correlations

#### Flight rate and proportion of time spent within 1m

```
##
##  Pearson's product-moment correlation
##
## data:  Robin2021$FlightRate and Robin2021$prop1m
## t = 2.2194, df = 40, p-value = 0.03219
## alternative hypothesis: true correlation is not equal to 0
## 95 percent confidence interval:
##  0.03023823 0.57698971
## sample estimates:
##       cor
## 0.3311264
```

#### Flight rate and closest approach(m)

```
##
##  Pearson's product-moment correlation
##
## data:  Robin2021$FlightRate and Robin2021$ClosestApproachM
## t = -2.3596, df = 40, p-value = 0.02326
## alternative hypothesis: true correlation is not equal to 0
## 95 percent confidence interval:
##  -0.59072113 -0.05103777
## sample estimates:
##       cor
## -0.349547
```

#### Closest approach(m) and proportion of time spent within 1m

```
##
##  Pearson's product-moment correlation
##
## data:  Robin2021$ClosestApproachM and Robin2021$prop1m
## t = -2.5647, df = 40, p-value = 0.01418
## alternative hypothesis: true correlation is not equal to 0
## 95 percent confidence interval:
##  -0.61004629 -0.08112558
## sample estimates:
##        cor
## -0.3757912
```

### Aggression PCA

```
## Principal Components Analysis
## Call: principal(r = PhysicalMeasures, nfactors = 1, residuals = FALSE,
##     rotate = "none", covar = FALSE)
## Standardized loadings (pattern matrix) based upon correlation matrix
##                              PC1   h2   u2 com
## Robin2021$FlightRate        0.65 0.42 0.58   1
## Robin2021$ClosestApproachM -0.85 0.73 0.27   1
## Robin2021$prop5m            0.83 0.68 0.32   1
##
##                 PC1
## SS loadings    1.83
## Proportion Var 0.61
##
## Mean item complexity =  1
## Test of the hypothesis that 1 component is sufficient.
##
## The root mean square of the residuals (RMSR) is  0.2
##  with the empirical chi square  9.75  with prob <  NA
##
## Fit based upon off diagonal values = 0.79
```

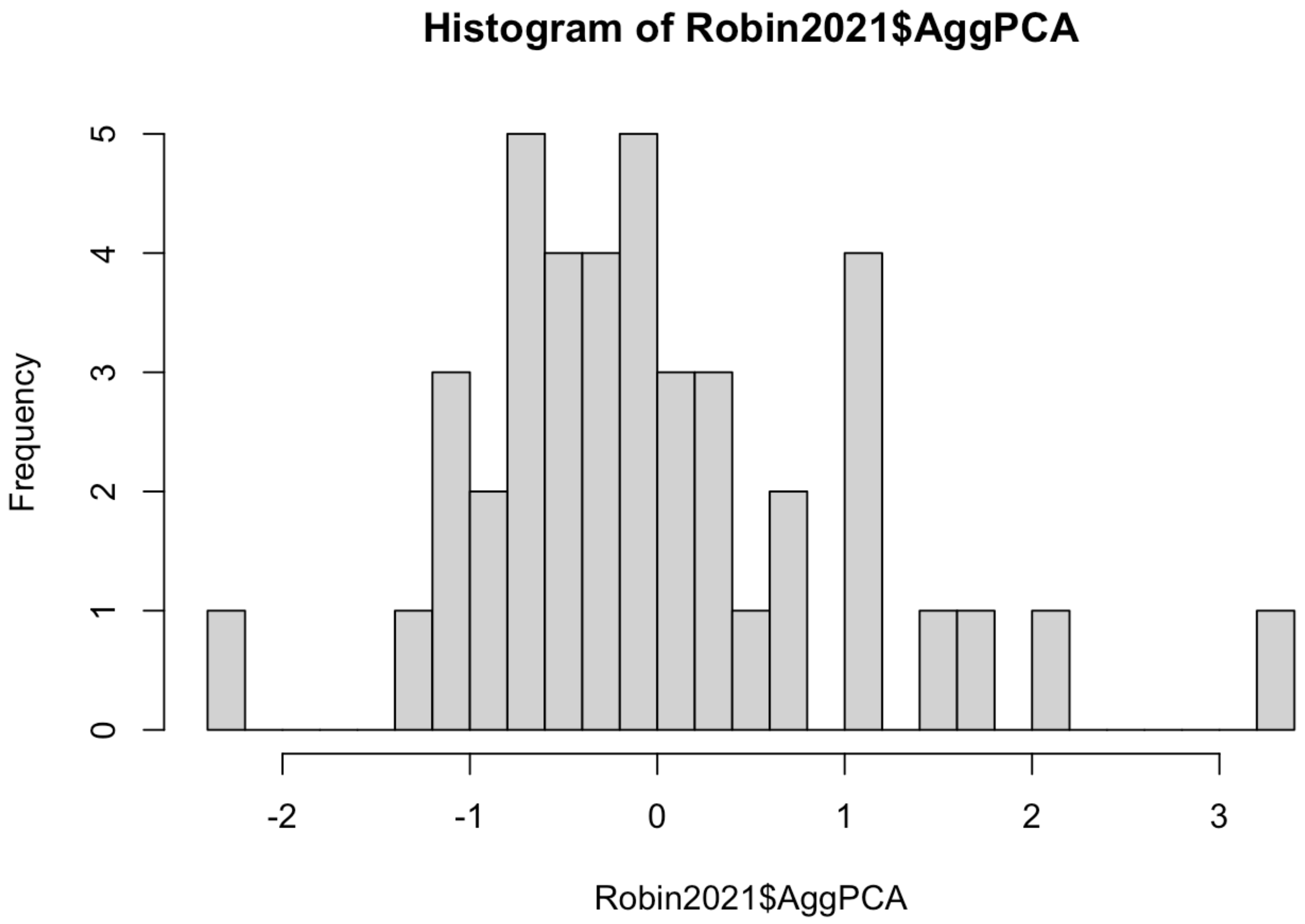


### Ambient Noise

#### Ambient Noise and Habitat correlation, LMM

```
## Warning: NAs introduced by coercion
```

```
## Linear mixed-effects model fit by REML
##   Data: Robin2021
##        AIC      BIC    logLik
##   204.1462 210.5899 -98.07312
##
## Random effects:
##  Formula: ~1 | MaleID
##         (Intercept) Residual
## StdDev:     5.27673 1.324208
##
## Fixed effects:  Average ~ Habitat
##                 Value Std.Error DF   t-value p-value
## (Intercept)  39.69781  1.550891 19 25.596782  0.0000
## HabitatUrban  8.96880  2.367907 19  3.787648  0.0012
##  Correlation:
##              (Intr)
## HabitatUrban -0.655
##
## Standardized Within-Group Residuals:
##         Min          Q1         Med          Q3         Max
## -1.31435938 -0.58807449 -0.05934325  0.59745834  1.62135675
##
## Number of Observations: 39
## Number of Groups: 21
```

```
##             numDF denDF   F-value p-value
## (Intercept)     1    19 1380.5883  <.0001
## Habitat         1    19   14.3463  0.0012
```

| | Ambient Noise | | |
|---|---|---|---|
| *Predictors* | *Estimates* | *CI* | *p* |
| (Intercept) | 39.70 | 36.45 – 42.94 | **<0.001** |
| Habitat Urban | 8.97 | 4.01 – 13.92 | **0.001** |
| **Random Effects** | | | |
| $\sigma^2$ | 1.75 | | |
| $\tau_{00}$ MaleID | 27.84 | | |
| $N_{MaleID}$ | 21 | | |
| Observations | 39 | | |
| Marginal $R^2$ / Conditional $R^2$ | 0.920 / NA | | |

### Ambient noise by habitat, Mean and SD

#### Urban

```
## [1] 48.98088
```

```
## [1] 7.071437
```

#### Rural

```
## [1] 39.94472
```

```
## [1] 3.570615
```

### Order effect

### Order effect on visual displays

```
## Generalized linear mixed model fit by maximum likelihood (Laplace
##   Approximation) [glmerMod]
##  Family: binomial  ( logit )
## Formula: VisualDisplay ~ Order + (1 | MaleID)
##    Data: Robin2021
##
##      AIC      BIC   logLik deviance df.resid
##     31.9     37.1    -12.9     25.9       39
##
## Scaled residuals:
##       Min        1Q    Median        3Q       Max
## -0.006653 -0.003376 -0.000002 -0.000002  0.096861
##
## Random effects:
##  Groups Name        Variance Std.Dev.
##  MaleID (Intercept) 3371     58.06
## Number of obs: 42, groups:  MaleID, 21
##
## Fixed effects:
##             Estimate Std. Error z value Pr(>|z|)
## (Intercept)  -11.344      3.358  -3.379 0.000729 ***
## Ordersecond  -15.323      6.221  -2.463 0.013778 *
## ---
## Signif. codes:  0 '***' 0.001 '**' 0.01 '*' 0.05 '.' 0.1 ' ' 1
##
## Correlation of Fixed Effects:
##             (Intr)
## Ordersecond 0.474
```

```
## Analysis of Variance Table
##       npar  Sum Sq Mean Sq F value
## Order    1 0.16273 0.16273  0.1627
```

### LMMs

#### Aggression, habitat and condition

```
## Linear mixed-effects model fit by REML
##   Data: Robin2021
##        AIC      BIC    logLik
##   105.9003 115.7259 -46.95017
##
## Random effects:
##  Formula: ~1 | MaleID
##         (Intercept)  Residual
## StdDev:   0.7549836 0.4650489
##
## Fixed effects:  AggPCA ~ Habitat * Condition
##                                 Value Std.Error DF   t-value p-value
## (Intercept)                -0.5826742 0.2559738 19 -2.276304  0.0346
## HabitatUrban                1.3208684 0.3910064 19  3.378125  0.0032
## Conditionnoise              0.3229697 0.1898554 19  1.701135  0.1052
## HabitatUrban:Conditionnoise -0.6761866 0.2900089 19 -2.331606  0.0309
##  Correlation:
##                             (Intr) HbttUr Cndtnn
## HabitatUrban                -0.655
## Conditionnoise              -0.371  0.243
## HabitatUrban:Conditionnoise  0.243 -0.371 -0.655
##
## Standardized Within-Group Residuals:
##         Min          Q1         Med          Q3         Max
## -1.68361774 -0.33760029  0.01751531  0.36962830  1.95791392
##
## Number of Observations: 42
## Number of Groups: 21
```

```
##                   numDF denDF  F-value p-value
## (Intercept)           1    19 0.000000  1.0000
## Habitat               1    19 7.324809  0.0140
## Condition             1    19 0.053435  0.8197
## Habitat:Condition     1    19 5.436387  0.0309
```

| | Aggression Score | | |
|---|---|---|---|
| *Predictors* | *Estimates* | *std. Error* | *p* |
| (Intercept) | -0.58 | 0.26 | **0.035** |
| Habitat Urban | 1.32 | 0.39 | **0.003** |
| Condition [noise] | 0.32 | 0.19 | 0.105 |
| Habitat Urban * Condition [noise] | -0.68 | 0.29 | **0.031** |

**Random Effects**

| | |
|---|---|
| $\sigma^2$ | 0.22 |
| $\tau_{00\ MaleID}$ | 0.57 |
| ICC | 0.72 |
| $N_{MaleID}$ | 21 |
| Observations | 42 |
| Marginal $R^2$ / Conditional $R^2$ | 0.257 / 0.795 |

### Urban Subset

```
## Linear mixed-effects model fit by REML
##   Data: Robin2021
##   Subset: Habitat == "Urban"
##        AIC      BIC    logLik
##   54.86982 57.96017 -23.43491
##
## Random effects:
##  Formula: ~1 | MaleID
##         (Intercept)  Residual
## StdDev:   0.8687837 0.6074828
##
## Fixed effects:  AggPCA ~ Condition
##                      Value Std.Error DF   t-value p-value
## (Intercept)      0.7381942 0.3533680  8  2.089024  0.0701
## Conditionnoise -0.3532169 0.2863701  8 -1.233428  0.2524
##  Correlation:
##                (Intr)
## Conditionnoise -0.405
##
## Standardized Within-Group Residuals:
##         Min          Q1         Med          Q3         Max
## -1.26451755 -0.53660150 -0.02836087  0.46799618  1.52319947
##
## Number of Observations: 18
## Number of Groups: 9
```

| | Aggression Score | | |
|---|---|---|---|
| *Predictors* | *Estimates* | *std. Error* | *p* |
| (Intercept) | 0.74 | 0.35 | 0.070 |
| Condition [noise] | -0.35 | 0.29 | 0.252 |

**Random Effects**

| | |
|---|---|
| $\sigma^2$ | 0.37 |
| $\tau_{00\ MaleID}$ | 0.75 |
| ICC | 0.83 |

| | |
|---|---|
| N $_{MaleID}$ | 21 |
| Observations | 18 |
| Marginal $R^2$ / Conditional $R^2$ | 0.015 / 0.829 |

### Song Rate, habitat and condition

```
## Linear mixed-effects model fit by REML
##   Data: Robin2021
##        AIC      BIC    logLik
##   194.9276 204.7531 -91.46378
##
## Random effects:
##  Formula: ~1 | MaleID
##         (Intercept) Residual
## StdDev:   0.9699014 2.185292
##
## Fixed effects:  SongRateTrial ~ Habitat * Condition
##                                 Value Std.Error DF   t-value p-value
## (Intercept)                  6.750000 0.6901819 19  9.780031  0.0000
## HabitatUrban                 0.509259 1.0542702 19  0.483044  0.6346
## Conditionnoise               2.277778 0.8921419 19  2.553156  0.0194
## HabitatUrban:Conditionnoise -1.018519 1.3627693 19 -0.747389  0.4640
##  Correlation:
##                             (Intr) HbttUr Cndtnn
## HabitatUrban                -0.655
## Conditionnoise              -0.646  0.423
## HabitatUrban:Conditionnoise  0.423 -0.646 -0.655
##
## Standardized Within-Group Residuals:
##        Min         Q1        Med         Q3        Max
## -3.1364985 -0.4378008  0.2796373  0.6017053  1.4510900
##
## Number of Observations: 42
## Number of Groups: 21
```

```
##                   numDF denDF  F-value p-value
## (Intercept)           1    19 392.6518  <.0001
## Habitat               1    19   0.0000  1.0000
## Condition             1    19   7.4543  0.0133
## Habitat:Condition     1    19   0.5586  0.4640
```

| | Song Rate | | |
|---|---|---|---|
| *Predictors* | *Estimates* | *std. Error* | *p* |
| (Intercept) | 6.75 | 0.69 | **<0.001** |
| Habitat Urban | 0.51 | 1.05 | 0.635 |
| Condition [noise] | 2.28 | 0.89 | **0.019** |

| | | | |
|---|---|---|---|
| Habitat Urban * Condition [noise] | -1.02 | 1.36 | 0.464 |

**Random Effects**

| | |
|---|---|
| $\sigma^2$ | 4.78 |
| $\tau_{00\ MaleID}$ | 0.94 |
| ICC | 0.16 |
| $N_{MaleID}$ | 21 |
| Observations | 42 |
| Marginal $R^2$ / Conditional $R^2$ | 0.140 / 0.282 |

### Song length, habitat and condition

```
## Linear mixed-effects model fit by REML
##   Data: Robin2021
##        AIC      BIC    logLik
##   65.71167 75.53719 -26.85584
##
## Random effects:
##  Formula: ~1 | MaleID
##         (Intercept) Residual
## StdDev:    0.428188 0.281638
##
## Fixed effects:  SongLengthAvg ~ Habitat * Condition
##                               Value Std.Error DF   t-value p-value
## (Intercept)               1.8149560 0.1479484 19 12.267489  0.0000
## HabitatUrban              0.0749190 0.2259950 19  0.331508  0.7439
## Conditionnoise            0.0672782 0.1149782 19  0.585139  0.5653
## HabitatUrban:Conditionnoise 0.0738884 0.1756321 19  0.420700  0.6787
##  Correlation:
##                           (Intr) HbttUr Cndtnn
## HabitatUrban              -0.655
## Conditionnoise            -0.389  0.254
## HabitatUrban:Conditionnoise  0.254 -0.389 -0.655
##
## Standardized Within-Group Residuals:
##          Min           Q1          Med           Q3          Max
## -1.728514316 -0.475640121  0.007346797  0.432370121  2.097382618
##
## Number of Observations: 42
## Number of Groups: 21
```

```
##                   numDF denDF  F-value p-value
## (Intercept)           1    19 338.7094  <.0001
## Habitat               1    19   0.2886  0.5974
## Condition             1    19   1.2960  0.2691
## Habitat:Condition     1    19   0.1770  0.6787
```

| | Average Song Length | | |
|---|---|---|---|
| *Predictors* | *Estimates* | *std. Error* | *p* |
| (Intercept) | 1.81 | 0.15 | **<0.001** |
| Habitat Urban | 0.07 | 0.23 | 0.744 |
| Condition [noise] | 0.07 | 0.11 | 0.565 |
| Habitat Urban * Condition [noise] | 0.07 | 0.18 | 0.679 |
| **Random Effects** | | | |
| $\sigma^2$ | 0.08 | | |
| $\tau_{00\ MaleID}$ | 0.18 | | |
| ICC | 0.70 | | |
| $N_{MaleID}$ | 21 | | |
| Observations | 42 | | |
| Marginal $R^2$ / Conditional $R^2$ | 0.022 / 0.705 | | |

#GLM

### Visual Signals, habitat and condition

```
##
## Call:
## glm(formula = VisualDisplay ~ Habitat * Condition, family = binomial,
##     data = Robin2021, subset = Order == "first")
##
## Deviance Residuals:
##      Min        1Q    Median        3Q       Max
## -0.75853  -0.60386  -0.60386   0.00013   1.89302
##
## Coefficients:
##                              Estimate Std. Error z value Pr(>|z|)
## (Intercept)                -1.609e+00  1.095e+00  -1.469    0.142
## HabitatUrban                2.018e+01  2.917e+03   0.007    0.994
## Conditionnoise              1.435e-15  1.549e+00   0.000    1.000
## HabitatUrban:Conditionnoise -1.966e+01  2.917e+03  -0.007    0.995
##
## (Dispersion parameter for binomial family taken to be 1)
##
##     Null deviance: 27.910  on 20  degrees of freedom
## Residual deviance: 15.312  on 17  degrees of freedom
## AIC: 23.312
##
## Number of Fisher Scoring iterations: 17
```

```
## Analysis of Deviance Table
##
## Model: binomial, link: logit
##
## Response: VisualDisplay
##
## Terms added sequentially (first to last)
##
##
##                   Df Deviance Resid. Df Resid. Dev
## NULL                                20     27.910
## Habitat            1   5.6395        19     22.271
## Condition          1   3.2297        18     19.041
## Habitat:Condition  1   3.7288        17     15.312
```

### Urban Subset

```
##
## Call:
## glm(formula = VisualDisplay ~ Condition, family = binomial, data = Robin2021,
##     subset = Order == "first" & Habitat == "Urban")
##
## Deviance Residuals:
##      Min        1Q    Median        3Q       Max
## -0.75853  -0.75853   0.00005   0.00005   1.66511
##
## Coefficients:
##                 Estimate Std. Error z value Pr(>|z|)
## (Intercept)        20.57    7929.26   0.003    0.998
## Conditionnoise    -21.66    7929.26  -0.003    0.998
##
## (Dispersion parameter for binomial family taken to be 1)
##
##     Null deviance: 11.4573  on 8  degrees of freedom
## Residual deviance:  4.4987  on 7  degrees of freedom
## AIC: 8.4987
##
## Number of Fisher Scoring iterations: 19
```

```
## Analysis of Deviance Table
##
## Model: binomial, link: logit
##
## Response: VisualDisplay
##
## Terms added sequentially (first to last)
##
##
##           Df Deviance Resid. Df Resid. Dev
## NULL                          8    11.4573
## Condition  1   6.9586         7     4.4987
```

### Plots

#### Visual signals, habitat and condition

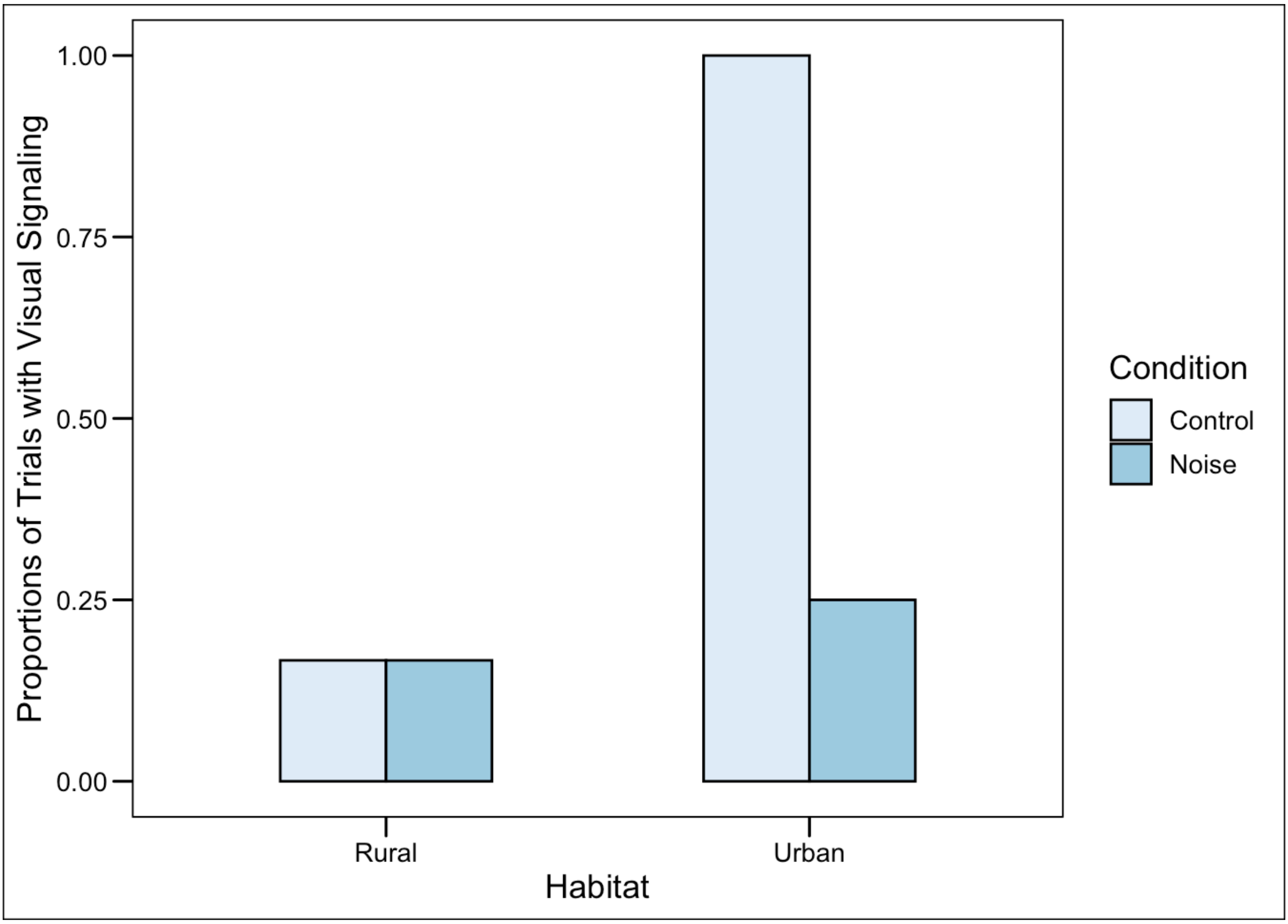


#### Song rate, habitat and condition

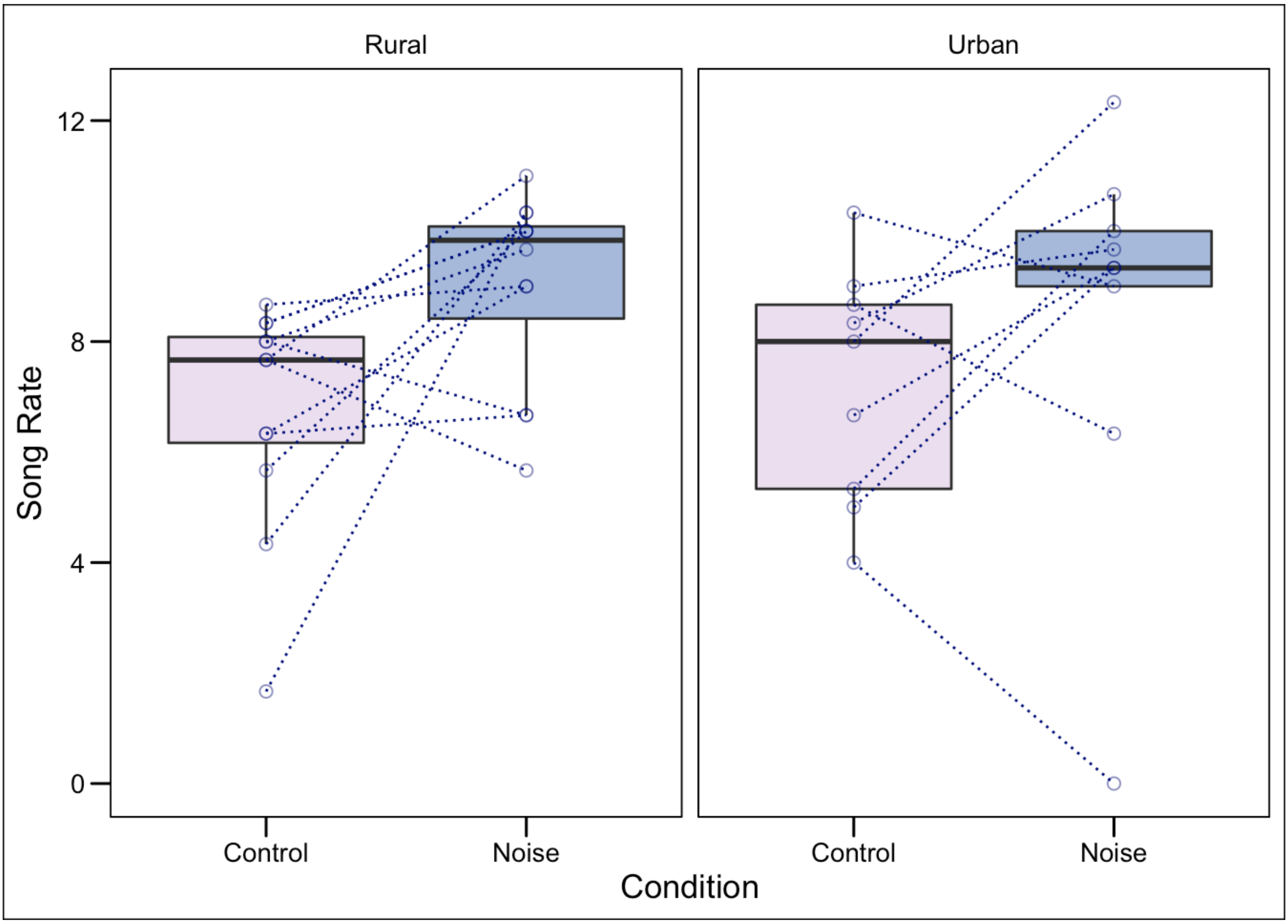


#### Aggression, habitat and condition

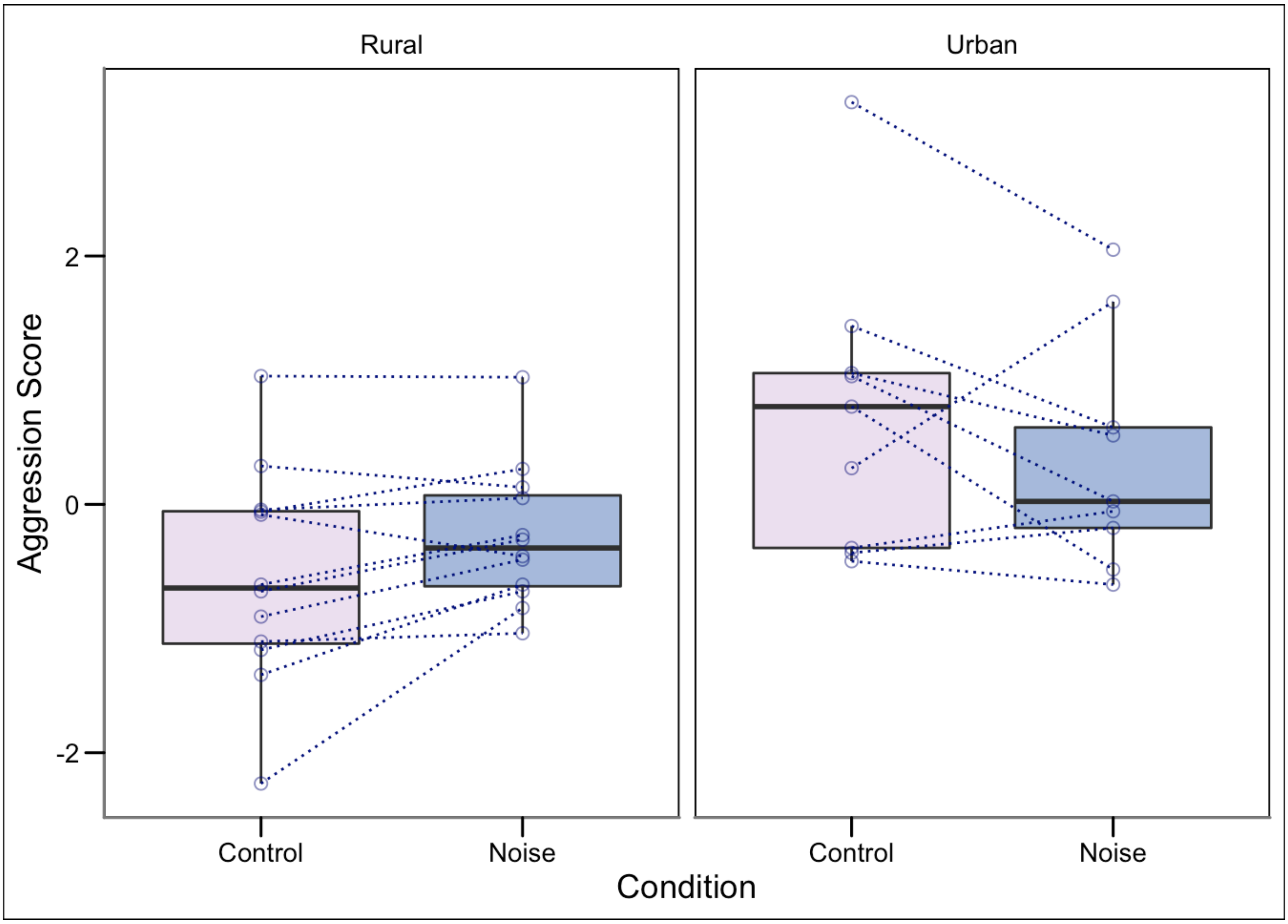
Rural
Urban
Aggression Score
2
0
-2
Control
Noise
Control
Noise
Condition
